# Potassium regulates axon-oligodendrocyte signaling and metabolic coupling in white matter

**DOI:** 10.1101/2022.11.08.515614

**Authors:** Zoe J. Looser, Luca Ravotto, Ramona B. Jung, Hauke B. Werner, Torben Ruhwedel, Wiebke Möbius, Dwight E. Bergles, L. Felipe Barros, Klaus-Armin Nave, Bruno Weber, Aiman S. Saab

## Abstract

The integrity of myelinated axons relies on homeostatic support from oligodendrocytes (OLs), which is essential for brain function. However, the mechanisms by which OLs detect axonal spiking and rapidly control axon-OL metabolic coupling are largely unknown. Here, we combine optic nerve electrophysiology and two-photon imaging to study activity-dependent calcium (Ca^2+^) dynamics in OLs and metabolite fluxes in myelinated axons. Both high-frequency axonal firing and extracellular potassium (K^+^) elevations trigger a fast Ca^2+^ response in OLs that is facilitated by barium-sensitive, inwardly rectifying K^+^ channels. Using OL-specific Kir4.1 knockout mice (Kir4.1 cKO) we now demonstrate that, in addition to being crucial for K^+^ clearance, oligodendroglial Kir4.1 regulates axonal energy metabolism and long-term axonal integrity. Before the manifestation of axonal damage, we observed reduced glucose transporter GLUT1 and monocarboxylate transporter MCT1 expression in myelin of young Kir4.1 cKO mice, suggesting early deficits in metabolite supply to axons. Strikingly, we found lower resting lactate levels and activity-induced lactate surges in optic nerve axons of young Kir4.1 cKO mice. Moreover, both axonal glucose uptake and consumption were hampered in the absence of oligodendroglial Kir4.1, uncovering a new role of OLs in regulating axonal glucose metabolism. Our findings reveal a novel model of axon-OL signaling and metabolic coupling in which OLs detect high-frequency axonal activity through K^+^ signaling, which is critical in adjusting the axon-OL metabolic unit and in preserving long-term axonal health.

## Main

Oligodendrocytes (OLs) produce and maintain the myelin sheaths around axons, enabling fast and economical electrical signal transmission between neurons. Axonal health is crucial for brain function, and axonal damage is a feature of aging and various neurological disorders^1, 2^. OLs have almost the same lifespan as neurons^3^ and apart from orchestrating axonal signaling speed, accumulating evidence reveals that OLs play an important role in preserving neural circuit functions and long-term neuronal integrity^4–8^. OLs are highly glycolytic and maintain axonal health by providing metabolic support to axons in form of lactate or pyruvate through monocarboxylate transporters^9–11^. The metabolic support machinery of OLs is likely adjusted by axonal activity, for example by regulating surface expression of glucose transporter 1 (GLUT1) in response to glutamatergic signaling^12^. Metabolite supply from OLs to axons is facilitated by cytosolic channels within the myelin sheaths^13, 14^ and a collapse of this myelinic channel system may lead to axonal damage^15^. Moreover, energy homeostasis is impaired in mice deficient of the myelin proteolipid protein (PLP)^16^, a mouse model of spastic paraplegia that develops severe axonal spheroids with age^17, 18^, possibly due to deficits in axonal transport^19^ and alterations in mitochondrial function^20^. OLs also carry out other homeostatic functions that are critical for axonal health, such as antioxidant support^21^ and K^+^ buffering^22, 23^.

However, the mechanisms governing the homeostatic interplay between axons and OLs are poorly explored. Here, we suspected that activity-driven K^+^ elevations may regulate metabolic coupling between OLs and axons. We addressed this question using our optic nerve electrophysiology and two-photon imaging technique, previously used to study axonal ATP dynamics^24, 25^. Imaging cytosolic Ca^2+^ dynamics in mature OLs revealed that high-frequency axonal activity is detected by OLs predominantly via increases in extracellular [K^+^] and Kir4.1 channel activation. Moreover, pharmacological inhibition of Kir4.1 impaired the recovery of axonal firing from high-frequency stimulation. Using OL-specific knockout mice (Kir4.1 cKO) we demonstrate that this is a specific function of oligodendroglial Kir4.1, implying that OLs are the primary cells involved in K^+^ clearance in white matter tracts. Kir4.1 cKO mice developed axonal pathology around 7-8 months of age. Significantly, we found reduced expression of GLUT1 and MCT1 in myelin purified from brains of 2-3 months old Kir4.1 cKO mice, suggesting reduced energy substrate delivery to myelinated axons prior to the onset of axonopathy. Using genetically encoded sensors, we studied axonal ATP, lactate and glucose dynamics in young Kir4.1 cKO mice, detecting lower lactate levels and activity-induced lactate surges. Axonal glucose uptake and consumption were also reduced, revealing for the first time that OLs regulate axonal glucose metabolism. These early metabolic deficits could affect axonal antioxidant capacity leading to late-onset axonal damage. Our findings teach that a rise in extracellular [K^+^] during fast axonal spiking stimulates OLs, and that oligodendroglial K^+^ homeostasis regulates axonal energy metabolism, function and survival.

## Results

### Axonal spiking evokes calcium response in oligodendrocytes

To investigate possible calcium (Ca^2+^) dynamics in mature oligodendrocytes (OLs) as a function of electrical activity we used crosses of *PLP-CreERT* mice^26^ with Ai96 mice that express the cytosolic Ca^2+^ indicator GCaMP6s in a Cre-dependent manner (RCL-GCaMP6s^RCL-GCaMP6s, 27,28^). We used 3-5 months old *PLP-CreERT;RCL-GCaMP6s* mice that were treated with tamoxifen at the age of 6-8 weeks (**Fig. 1a**), and confirmed by immunohistochemistry that GCaMP6s expression was restricted to mature (CC1-immunopositive) OLs (**Fig. 1b**). We studied the optic nerve, a myelinated white matter tract model ideal for simultaneous recordings of compound action potentials (CAPs) and two-photon sensor imaging (**Fig. 1c**)^24^. To test if mature OLs sense axonal spiking activity, optic nerves were stimulated for 30 s at 10, 25 or 50 Hz. Before and after the high-frequency stimulation period, nerves received electrical pulses every 2.5 s (frequency of 0.4 Hz) to monitor the CAP changes in parallel to Ca^2+^ imaging in OLs. The peak amplitude of the CAP decreased during high frequency stimulation (**Fig. 1d**), as previously reported^24^. Importantly, we found that axonal stimulation triggered a bi-phasic Ca^2+^ response in OL somas consisting of an initial rise in Ca^2+^ levels during stimulation followed by a transient undershoot after stimulation (**Fig. 1e, Movie S1**). The electrically-evoked Ca^2+^ response was significantly larger at higher frequencies (**Fig. 1e**). Application of tetrodotoxin (TTX, 1 µM) abrogated the stimulus-induced Ca^2+^ surge in OLs (**Fig. 1f**), confirming the necessary role of action potentials. Moreover, removal of extracellular Ca^2+^ strongly diminished the elicited OL Ca^2+^ response (**Fig. 1g**), suggesting a mechanism involving Ca^2+^ influx.

**Figure 1.**
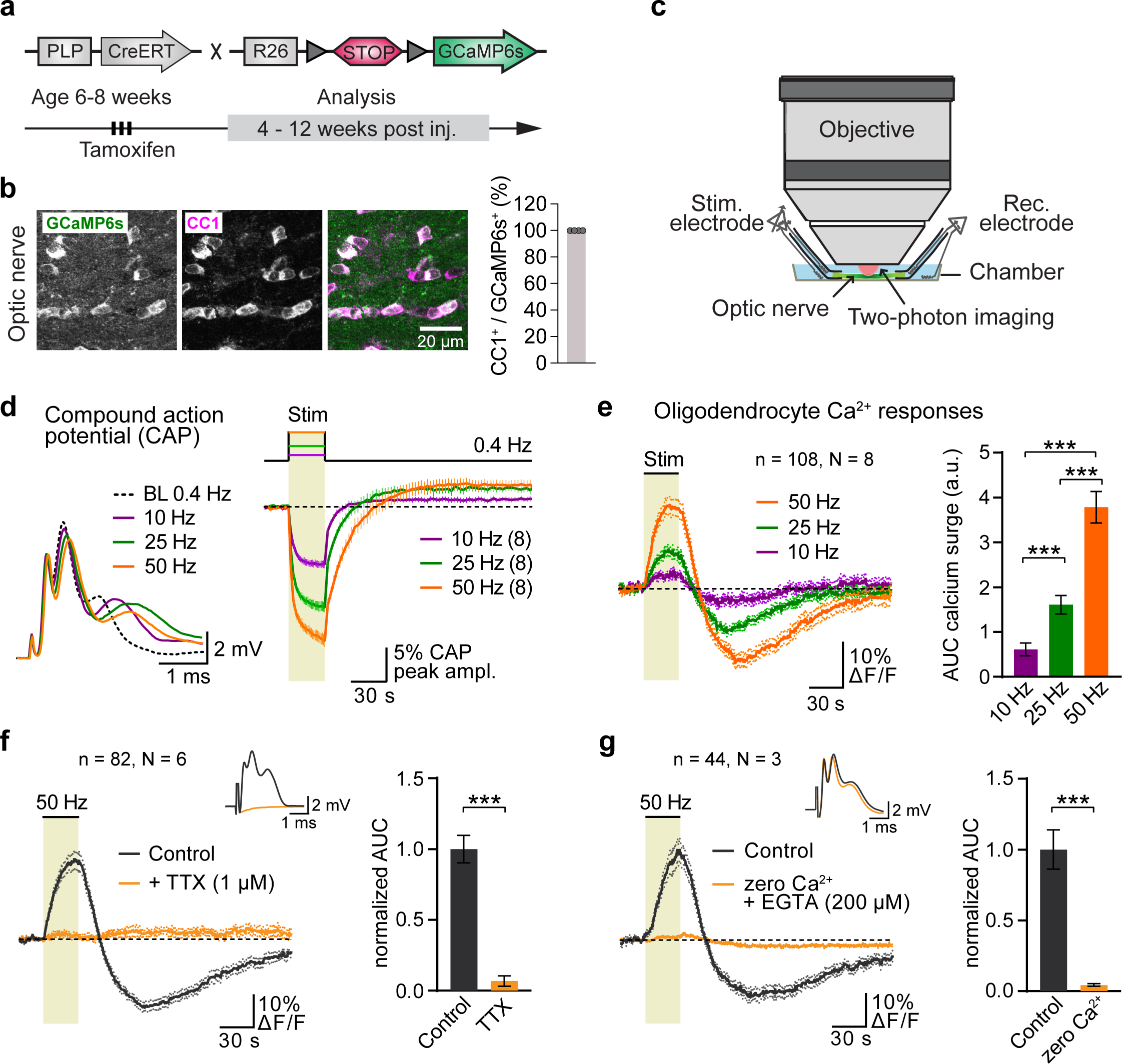
Optic nerve axonal activity-induced Ca^2+^ dynamics in oligodendrocytes (OLs) **a,** Generation of *PLP-CreERT;RCL-GCaMP6s* mice. Tamoxifen treatment for 3 consecutive days at the age of 6 to 8 weeks and experiments were conducted 4 to 12 weeks post injections. **b**, Immunohistochemical analysis of GCaMP6s (with α-GFP antibody, green) and CC1 (magenta), a mature OL marker, in optic nerve sections confirms OL-specific expression of GCaMP6s (quantification of a total 12 sections from 4 mice). Scalebar 20 µm. **c**, Acute optic nerve preparation for combined electrophysiology and two-photon imaging. **d**, Example traces of the three-peak shaped CAP response at 0.4 Hz baseline (BL) and at the end of a 30 s stimulation train with 10, 25 or 50 Hz. Average time course of relative changes in peak 2 amplitude upon higher frequency stimulations. Changes shown relative to the 0.4 Hz BL recordings (N = 8 optic nerves). **e**, Average OL Ca^2+^ responses at different frequency axonal stimulations. Bar graph depicts quantification of the area under the curve (AUC) of individual OL soma (n = 108, N = 8 optic nerves) responses during the 30 s stimulation period. Stimulations with higher frequencies evoked a larger Ca^2+^ surge in OLs (50 Hz vs. 25 Hz, p < 0.0001; 50 Hz vs. 10 Hz, p <0.0001; 25 Hz vs. 10 Hz, p < 0.0001; one-way ANOVA with Tukey’s multiple comparisons test). **f**, TTX (1 μM) abolished the 50 Hz stimulation-induced OL Ca^2+^ response. Insert: CAP activity diminished by TTX. Normalized AUCs of evoked responses from individual OLs before and after TTX application (n = 82, N = 6), showing a reduction by 93 ± 9% (p < 0.0001, paired t-test). **g**, Removal of extracellular Ca^2+^ (plus addition of 200 μM EGTA) diminished the stimulation-evoked OL Ca^2+^ response by 96 ± 14% (n = 44, N = 3, p < 0.0001, paired t-test). Data are represented as means ± SEM.

OLs express NMDA receptors that regulate oligodendroglial glucose import and metabolic support to axons^12^. Furthermore, they mediate a Ca^2+^ rise in myelin upon electrical stimulation of axons^29^ and in response to chemical ischemia^30^. AMPA receptors were also suggested to play a role in activity-mediated Ca^2+^ dynamics in myelin^29^. We therefore tested if glutamatergic signaling contributes to the stimulus-evoked OL Ca^2+^ response (**Extended Data Fig. 1a-d)**. No difference in the OL Ca^2+^ response was detected when AMPA receptors were blocked with NBQX (50 µM; **Extended Data Fig. 1b**), suggesting that AMPA receptors are not involved. However, additional inhibition of NMDA receptors with 7-CKA (100 µM) and D-AP5 (100 µM), to block both glycinergic and glutamatergic NMDA receptor binding sites, led to a reduction of about 20% in the evoked OL Ca^2+^ response (**Extended Data Fig. 1d**). We next examined purinergic signaling, given that ATP has also been implicated as a signaling molecule in white matter and OLs express P2X/P2Y receptors^31–34^. The broad-spectrum, nonselective P2X/P2Y receptor antagonists PPADS (50 µM) and Suramin (50 µM) reduced the stimulus-evoked Ca^2+^ response by 20% (**Extended Data Fig. 1e-h**). This suggests that purinergic signaling, like glutamatergic signaling, has a minor contribution to the OL Ca^2+^ response.

Axonal action potentials cause an increase of extracellular [K^+^] in white matter tracts^22, 35, 36^, and we tested if K^+^ is the responsible signal to stimulate OLs. Indeed, we found that a transient bath application of K^+^ ([K^+^]_bath_) induced a Ca^2+^ response, which was more pronounced at higher K^+^ concentrations (**Fig. 2a**). Importantly, we ruled out the possibility that raising [K^+^]_bath_ may indirectly stimulate axonal firing and possible neurotransmitter release, since in the presence of TTX, OLs showed the same [K^+^]_bath_-evoked Ca^2+^ response (**Fig. 2b**). OLs express Kir4.1^22, 23, 37^ that has been implicated in regulating extracellular K^+^ homeostasis and excitability of neurons^22^. Moreover, K^+^ uptake through Kir channels during the buildup of extracellular K^+^ upon axonal spiking facilitates OL depolarization^22, 37–39^. Strikingly, we found that application of 100 µM barium (Ba^2+^), which blocks Kir4.1 channels, reduced the evoked OL Ca^2+^ response by 80% and in reversible fashion (**Fig. 2c**). Additionally, Ba^2+^ inhibited the [K^+^]_bath_-evoked Ca^2+^ response in OLs (**Fig. 2d**), implying that depolarization of OLs facilitated by Ba^2+^-sensitive Kir channels participates in the Ca^2+^ surge.

**Figure 2.**
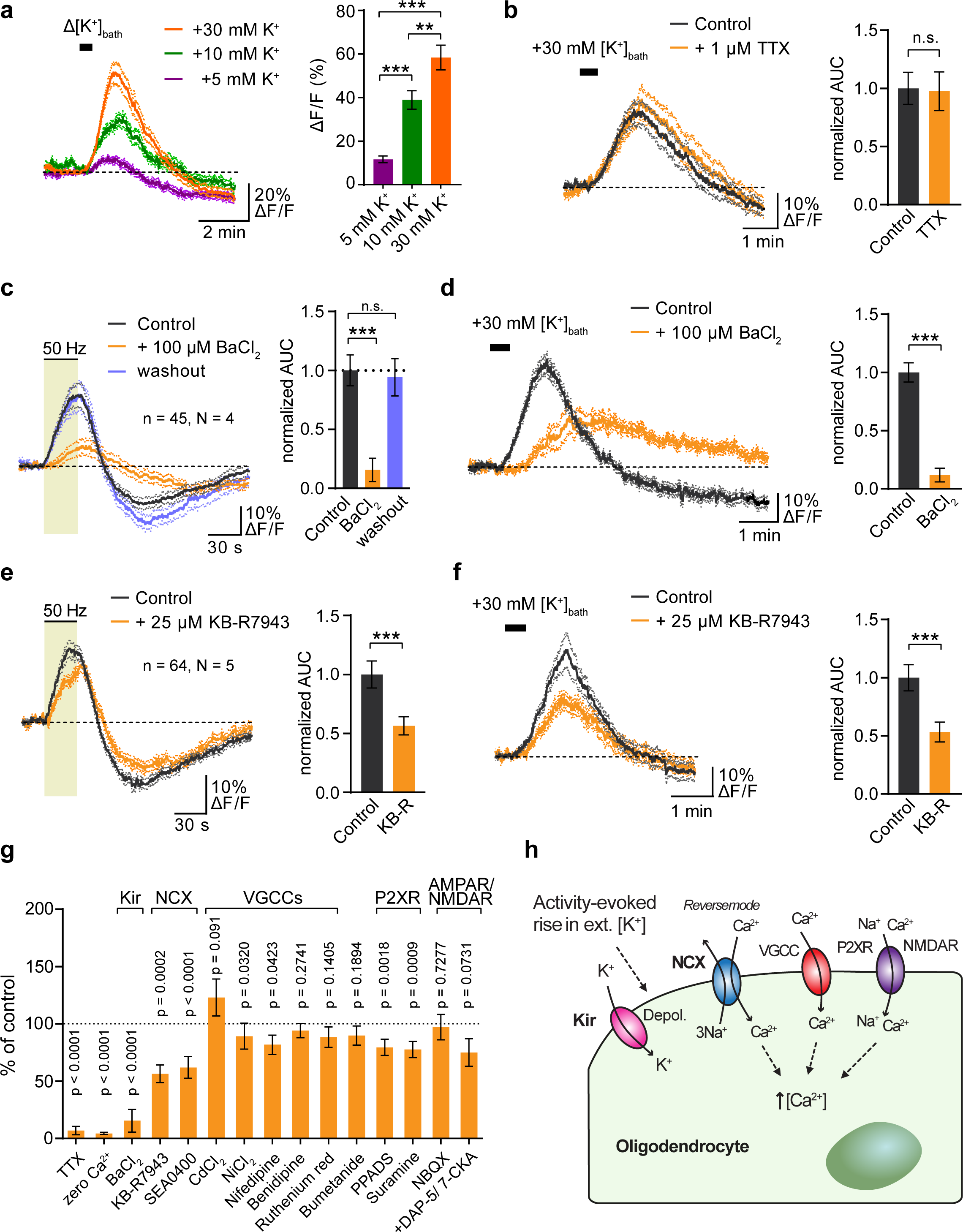
Kir channel-mediated mechanism underlying stimulus-evoked Ca^2+^ rise in OLs. **a,** Ca^2+^ levels in OLs increase to transiently raising extracellular [K^+^] by 5, 10 and 30 mM via 30 s bath application (Δ[K^+^]_bath_). Depicted are average Ca^2+^ traces of individual OLs and the quantification of Δ[K^+^]_bath_-evoked signal amplitudes (5 mM vs. 10 mM, p = 0.0048; 5 mM vs. 30 mM, p = <0.0001, 10 mM vs. 30 mM, p = <0.0001; n = 35-57 cells, N = 4-5 optic nerves; one way ANOVA with Tukey’s multiple comparisons test). **b**, The K^+^-evoked Ca^2+^ response in OLs is not due to secondary axonal spiking activity, as revealed by comparable K^+^-evoked Ca^2+^ surges in the presence of 1 µM TTX (n = 72, N = 5; p = 0.8144, paired t-test). **c**, Barium (Ba^2+^, 100 µM) reversibly inhibited the 50 Hz stimulation-induced OL Ca^2+^ surge by 84 ± 10% (n = 45, N = 4; p <0.0001, paired t-test). **d**, The K^+^-evoked Ca^2+^ rise in OLs was also reduced with Ba^2+^ by 88 ± 9% (n = 47, N = 3; p <0.0001, paired t-test). AUC was measured within one min of +30 mM [K^+^]_bath_-induced activation. **e**, The reverse-mode NCX blocker KB-R7943 (25 μM) reduced the 50 Hz stimulation-evoked Ca^2+^ rise by 44 ± 11% (n = 64, N = 5; paired t-test, p= 0.0002) as well as **f**, the K^+^-evoked response by 47 ± 8% (n = 52, N = 3; paired t-test, p = <0.0001). **g**, Summary of drugs tested and their inhibitory effects on the 50 Hz stimulation-evoked Ca^2+^ rise (see also Extended Data Figures 1, 3 and 4). **h**, Working scheme of axonal activity-mediated OL Ca^2+^ activation. A rise in [K^+^]_ext_ during high-frequency axonal activity depolarizes OLs via Kir channels, which facilitates Ca^2+^ entry through reverse-mode activation of NCX, whereas VGCCs, P2XR and NMDARs revealed minor contributions. Data are represented as means ± SEM.

Next, we tested if Ba^2+^ also affects axonal Ca^2+^ dynamics or whether the Ba^2+^-mediated inhibition of the stimulus-evoked Ca^2+^ response is specific to OLs. To study Ca^2+^ dynamics in optic nerve axons, we performed intravitreal delivery of adeno associated virus (AAV) for the expression of Cre in retinal ganglion cells of *RCL-GCaMP6s* mice, followed by two-photon imaging of GCaMP6s-expressing axons 3-6 weeks after AAV injection (**Extended Data Fig. 2a**). As expected, electrical stimulation elicited a strong Ca^2+^ rise in axons (**Extended Data Fig. 2b, Movie S2**), which was significantly larger at higher frequencies (**Extended Data Fig. 2c**). Notably, however, 100 µM Ba^2+^ did not affect the stimulus-evoked Ca^2+^ surge in axons (**Extended Data Fig. 2d**), emphasizing that the inhibition of the OL Ca^2+^ response with Ba^2+^ (**Fig. 2c**) was not due to an unspecific inhibition of axonal Ca^2+^ signaling. Yet, inhibition of glial Kir4.1 with Ba^2+^ had a strong impact on the recovery of axonal firing following 50 Hz stimulation as seen from the marked slower recovery of the CAP peak amplitude (**Extended Data Fig. 2e, f**).

Depolarization of OLs could activate voltage-gated Ca^2+^ channels (VGGCs) so we tested various VGGC blockers including cadmium (Cd^2+^), nickel (Ni^2+^), nifedipine, benidipine and ruthenium red (RuR, **Extended Data Fig. 3**), of which only Ni^2+^ and nifedipine revealed minimal effects (**Extended Data Fig. 3d, f)**. On the other hand, Cd^2+^, which did not affect the OL Ca^2+^ surge (**Extended Data Fig. 3b**), clearly reduced the stimulus-evoked axonal Ca^2+^ response (**Extended Data Fig. 2g, h**). This shows that the increase in axonal Ca^2+^ during electrical activity is mediated by activation of VGCCs, which is not a feature of the OL Ca^2+^ response. Moreover, the slower recovery of CAP conductance after high-frequency electrical activity was specific to Ba^2+^ (**Extended Data Fig. 2**), and was not affected by Cd^2+^ (**Extended Data Fig. 2i, j**).

**Figure 3.**
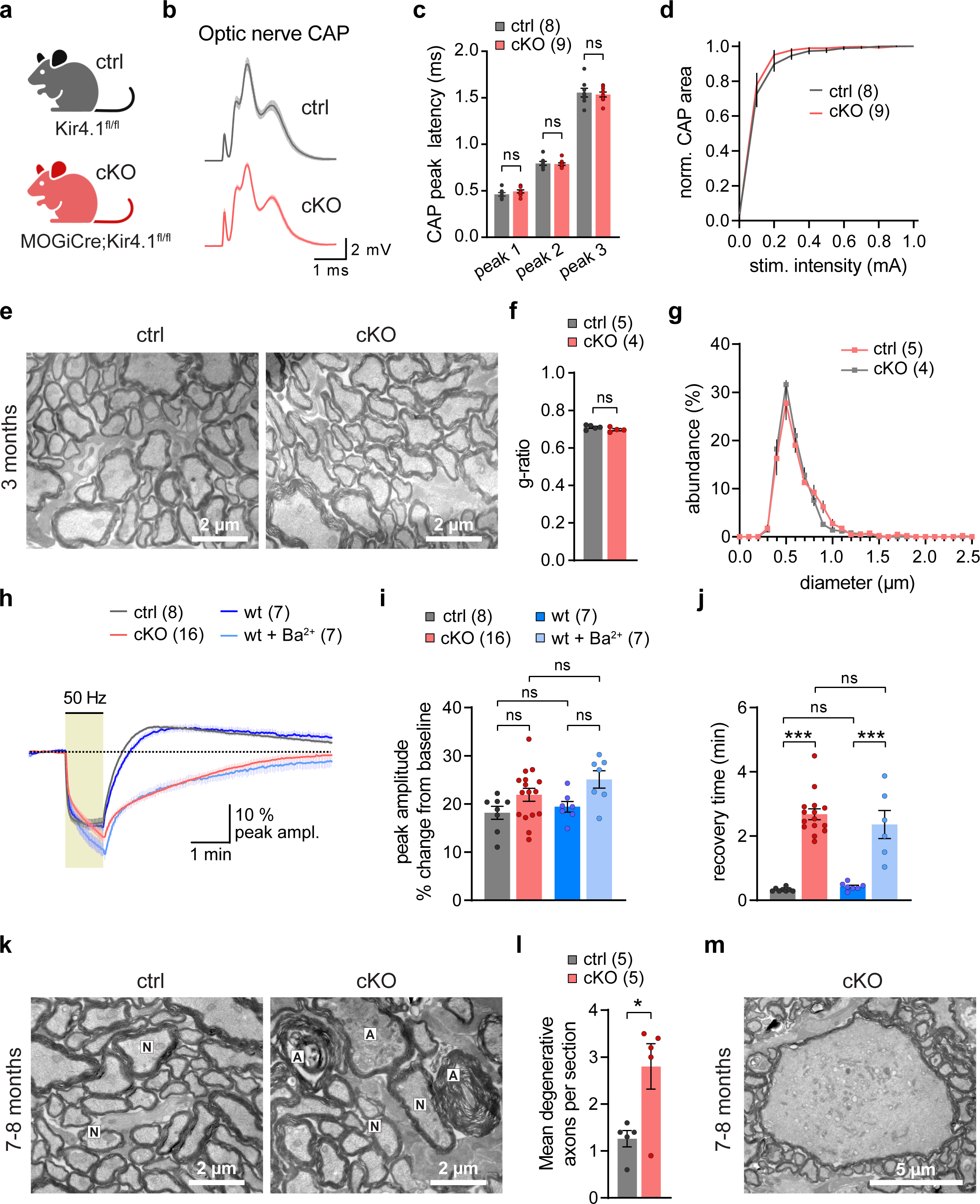
Oligodendroglial Kir4.1 is critical for white matter K^+^ clearance and long-term axonal integrity. **a,** OL-specific Kir4.1 knockout mice (*Kir4.1^fl/fl^;MOGiCre*, termed cKO) and littermate controls (*Kir4.1^fl/fl^*, termed ctrl). **b**, Average CAP response of optic nerves from ctrl (n = 8) and cKO (n = 9). **c**, No differences in CAP peak latencies between the genotypes (p = 07637 for peak 1, p = 0.9958 for peak 2, p = 0.9265 for peak 3, one-way ANOVA with Holm-Šídák’s multiple comparisons test). **d**, No difference in the stimulus-response relationships (*F_interaction_* (10, 150) = 0.4445, p = 0.9224, two-way ANOVA). CAP area from each stimulus intensity was normalized to the max stimulation of 1 mA for each nerve. **e-g**, Electron microscopic inspection of optic nerves from 3 months old cKO and ctrl mice (**e**) revealed no difference in myelin sheath thickness determined by (**f**) g-ratio analysis (n = 4-5 mice, p = 0.1584, unpaired t-test) and (**g**) normal axon size distribution of myelinated axons (*F_interaction_* (25, 175) = 1.028, p = 0.4333, two-way ANOVA). **h**, Averaged changes (in % from baseline) of CAP peak amplitude upon 1 min 50 Hz stimulation of optic nerves from cKO and ctrl as well as from wildtype (wt) nerves and wt treated with 100 µM Ba^2+^ (wt+Ba^2+^). Note that in recordings from cKO and wt+Ba^2+^ the recovery of the CAP peak amplitude following 50 Hz stimulation is similarly slower compared to ctrl and wt. **i**, Analysis of the CAP peak amplitude drop from baseline at the end of the 50 Hz stimulation period from recordings in (**h**). The stimulus-evoked reduction in amplitude was comparable between the groups (ctrl vs cKO, p = 0.2486; wt vs wt+Ba^2+^, p = 0.1040; ctrl vs wt, p = 0.9753; cKO vs wt+Ba^2+^, p = 0.4418; one-way ANOVA with Holm-Šídák’s multiple comparisons test). **j**, Analysis of the CAP peak recovery time following 50 Hz stimulation. Recovery is profoundly slower in nerves from cKO and wt+Ba^2+^ compared to ctrl and wt, respectively (ctrl vs cKO, p < 0.0001; wt vs wt+Ba^2+^, p < 0.0001; ctrl vs wt, p = 0.9970; cKO vs wt+Ba^2+^, p = 0.7362; one-way ANOVA with Holm-Šídák’s multiple comparisons test). **k-m**, Electron microscopic inspection of optic nerves from 7-8 months old cKO and ctrl mice (**k**) revealed a significant increase in ultrastructural features of axonal injury and degeneration in cKO (**l**) (n = 5 mice with 10 randomly taken images, each covering 286 µm^2^, p = 0.0169, unpaired t-test) and giant axonal swellings were unique to cKO nerves at that age (**m**). Data are represented as means ± SEM.

OLs express Na^+^/Ca^2+^ exchangers (NCX)^40–42^, and NCX are sensitive to membrane depolarization that may operate in reverse mode allowing Ca^2+^ entry^42, 43^. Inhibition of the sodium pumps with 500 µM ouabain caused a Ca^2+^ surge in OLs, which was reduced by blocking the reverse-mode activity of the NCX with 25 µM KB-R7943 (**Extended Data Fig. 4a, b**), confirming that functional NCX are expressed by optic nerve OLs. We then tested if activation of NCX contributes to the stimulus-evoked Ca^2+^ response. Indeed, 25 µM KB-R7943 reduced the OL Ca^2+^ response by 45% (**Fig. 2e**). We observed a similar inhibition of the evoked OL Ca^2+^ response using 10 µM SEA0400, another NCX blocker (**Extended Data Fig. 4c, d**). Additionally, we revealed that KB-R7943 also decreased the [K^+^]_bath_-evoked Ca^2+^ response in OLs (**Fig. 2f**). These results indicate that K^+^-mediated depolarization of OLs causes Ca^2+^ entry through reverse-mode activation of NCX.

**Figure 4.**
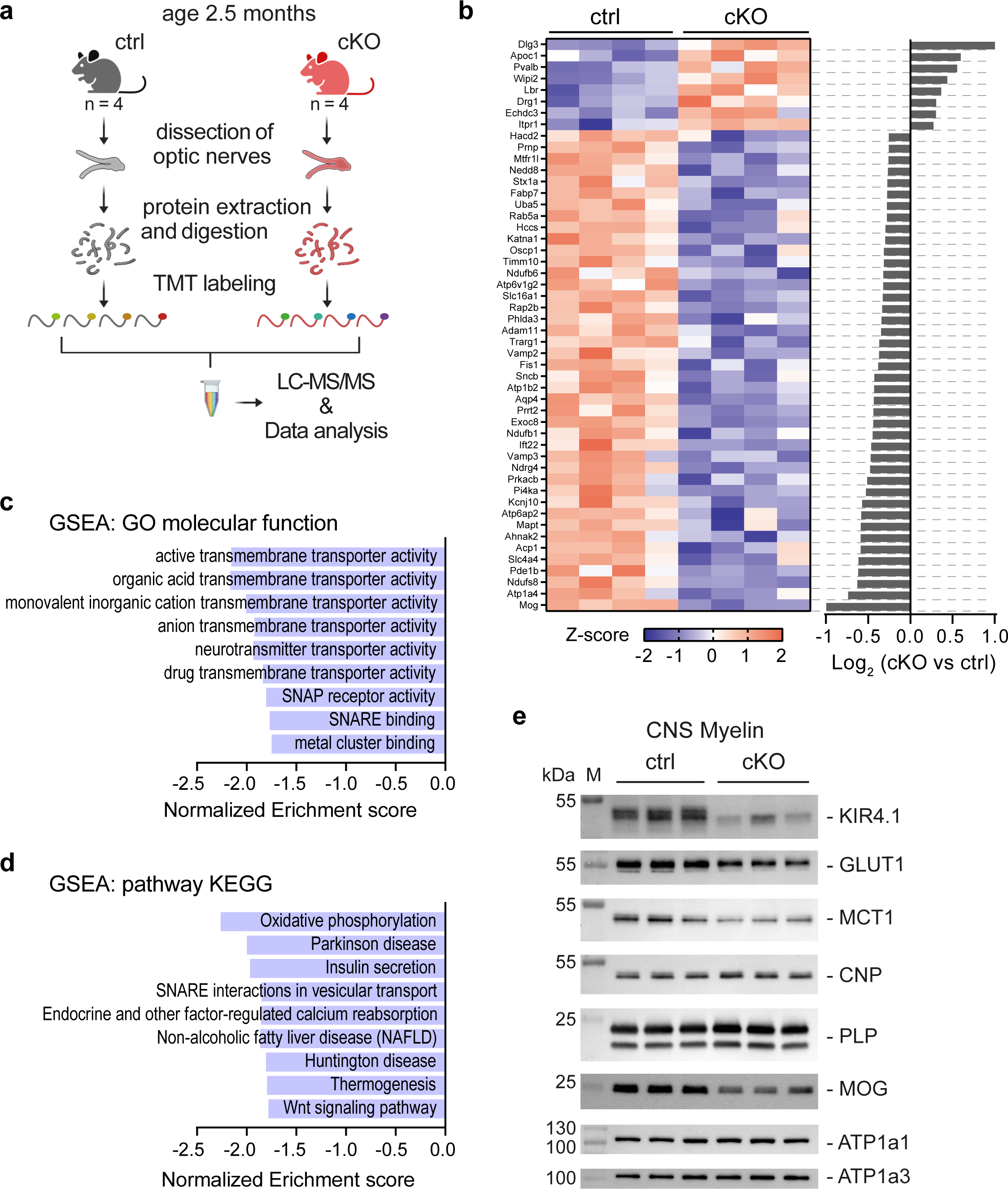
Reduced MCT1 and GLUT1 levels in CNS myelin of Kir4.1 cKO mice. **a,** Optic nerves were dissected from 2.5 months old cKO (n = 4) and ctrl (n =4) mice. Following protein extraction and digestion, proteins were labeled with tandem mass tags (TMT) and then pooled for liquid chromatography-tandem mass spectrometry (LC-MS/MS). Scheme created with BioRender. **b**, Heatmap showing relative up- (red) or downregulation (blue) of proteins ranked by log2 fold changes between cKO and ctrl mice (right bar graph). Depicted are the gene names of the top 50 differentially regulated proteins obtained from sorting identified proteins by false discovery rate (FDR) and restricted to >0.25 log2 fold changes between the genotypes. Row *Z*-scores were calculated using normalized protein intensities. **c** and **d**, Gene enrichment analyses (GSEAs) for categories (**c**) GO molecular function and (**d**) pathway KEGG with an FDR < 0.05. Note the overall decrease in pathways related to transmembrane transporter activity, vesicular transport and energy metabolism. WebGestalt.org was employed for analysis using as input list all identified proteins ranked by log2 fold changes between cKO and ctrl mice. **e**, Immunoblot analysis of Kir4.1, MCT1 and GLUT1 in myelin biochemically purified from the brains of 2.5 months old ctrl (n = 3) and cKO mice (n = 3). Compared to ctrl, the abundance of Kir4.1 was reduced by 81 ± 13% (p = 0.0038), of GLUT1 was reduced by 44 ± 8% (p = 0.0044) and of MCT1 was reduced by 50 ± 13% (p = 0.0179, unpaired t-tests). Known myelin proteins PLP, CNP, MOG, ATP1a1 and ATP1a3 are detected as markers.

Recent reports showed that Na^+^/K^+^/Cl^−^ cotransporter (NKCC1) is expressed in developing OLs^44, 45^ and may play a role in the volume regulation of the axon-facing inner tongue^44^. We tested if Na^+^/K^+^/Cl^−^ cotransporters are involved in the evoked OL Ca^2+^ surge using Bumetanide, a specific inhibitor of NKCC1. However, we found that 50 µM Bumetanide had no effect on the stimulus-induced Ca^2+^ rise in OLs (**Extended Data Fig. 4e, f**), ruling out that possible NKCC1-mediated volume changes in adult OLs could be a contributing factor during high-frequency axonal firing.

In summary, our pharmacological results (summarized in **Fig. 2g**) suggest that OLs detect high-frequency axonal activity via elevations in extracellular K^+^ concentrations that depolarize OLs via Kir channels, promoting Ca^2+^ entry chiefly through reverse-mode activation of NCX (**Fig. 2h**).

### Oligodendroglial Kir4.1 regulates metabolite transporter abundance in CNS myelin

We discovered that in the presence of Ba^2+^, the recovery kinetics of axonal firing after high frequency activity were significantly reduced (**Extended Data Fig. 2f**), suggesting that K^+^ clearance was impaired when Kir channels were blocked. A similar finding was reported in mice lacking Kir4.1 specifically from OLs^22^. We thus tested whether the Ba^2+^-mediated impact on CAP peak recovery was primarily attributable to suppression of oligodendroglial Kir4.1, or if astrocytes which also express Kir4.1^37, 46^ may also be implicated in K^+^ clearance in white matter^36^. To address this, we made use of the previously reported^22^ OL-specific Kir4.1 knockout mice (*Kir4.1^fl/fl^;MOGiCre*, hereafter termed cKO, **Fig. 3a**). First, we inspected the CAP properties of optic nerves from 2-3 months old Kir4.1 cKO mice and littermate controls (Kir4.1^fl/fl^, **Fig. 3b-d**). We found no differences in CAP peak latencies (**Fig. 3c**) and nerve excitability (**Fig. 3d**) between the genotypes, suggesting that myelination of optic nerve axons is not considerably perturbed in the absence of oligodendroglial Kir4.1. Accordingly, electron microscopic inspection (**Fig. 3e**) revealed no overt changes in myelin sheath thickness (**Fig. 3f**) and diameter distribution of myelinated axons (**Fig. 3g**). At this age, we also did not observe signs of axonal damage (**Fig. 3e**) or general neuroinflammation (**Extended Data Fig. 5a, b**). We then examined whether K^+^ clearance was perturbed in Kir4.1 cKO mice by measuring axonal conduction properties after high-frequency optic nerve stimulations (**Fig. 3h-i**). Indeed, following 50 Hz stimulation the recovery kinetics of the CAP peak amplitude were significantly slower in Kir4.1 cKO nerves compared to littermate controls (**Fig. 3h, j**), confirming earlier results^22^. Not only CAP peak amplitude but also the recovery kinetics of the CAP peak latency were significantly slower in cKO (**Extended Data Fig. 6a-d**). We then performed similar CAP recordings using optic nerves from wildtype mice in the presence of 100 µM Ba^2+^ and compared the conduction kinetics to Kir4.1 cKO results. Strikingly, wildtype nerves treated with Ba^2+^ showed the same reduced recovery kinetics of axonal firing as Kir4.1 cKO nerves (**Fig. 3h-j**). These results imply that oligodendroglial Kir4.1 is primarily involved in K^+^ buffering in the adult white matter with little to no contribution of astrocytic Kir4.1 or other Ba^2+^-sensitive Kir channels. The deficits in the CAP recovery kinetics after electrical activity were also visible at 25 Hz and 10 Hz stimulation frequencies (**Extended Data Fig. 6e-p**), demonstrating that oligodendroglial Kir4.1 also governs activity-dependent K^+^ clearance at lower frequency.

**Figure 5.**
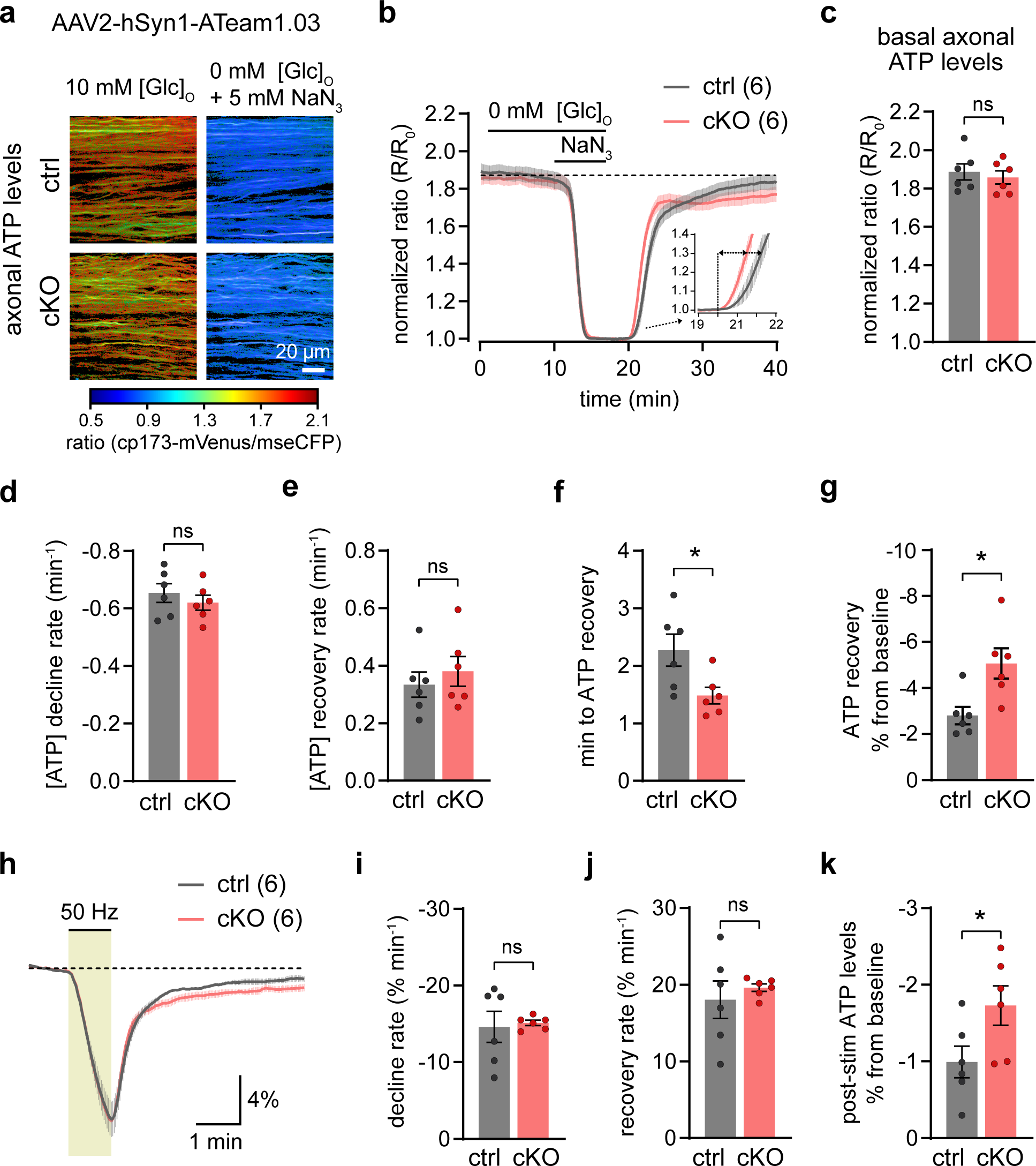
Axonal ATP dynamics slightly altered in young Kir4.1 cKO mice. **a,** ATP FRET sensor (ATeam1.03) expression in optic nerve axons via intravitreal AAV delivery. Depicted are example color-coded ratio images from acute optic nerves of ctrl (top) and cKO (bottom) in ACSF containing 10 mM glucose (Glc) and after glucose deprivation (GD) plus mitochondrial inhibition (MI) with 5 mM NaN3. Warm and cold colors indicate high and low ratios or ATP levels, respectively, and cp173-mVenus and mseCFP are FRET acceptor and donor, respectively. Scalebar 20 µm. **b**, Time course of axonal ATP level changes from optic nerves of ∼3 months old cKO and ctrl mice transiently challenged with GD+MI, FRET ratio (R) normalized to the minimum ATP level (R0) achieved by GD+MI. Inset depicts initial ATP recovery dynamics following reperfusion with 10 mM glucose. **c**, Basal axonal ATP levels obtained from (**b**) are comparable between the genotypes (n = 6, p = 0.61, unpaired t-test). **d**, No difference in the ATP decline rate upon GD+MI from (**b**) between the genotypes (n = 6, p = 0.44, unpaired t-test). **e**, Similar initial ATP recovery rates after GD+MI between the genotypes (n = 6, p = 0.51, unpaired t-test). **f**, Onset of ATP recovery (see inset in **b**) differs between ctrl and cKO nerves (n = 6, p = 0.03, unpaired t-test) **g**, Lower axonal ATP level recovery in cKO nerves following GD+MI compared to ctrl (n = 6, p = 0.0135, unpaired t-test). **h**, Axonal ATP level changes (in %) in response to 50 Hz stimulation. FRET ratios normalized to baseline before stimulation. **i** and **j**, Similar ATP level decline rates (**i**) during 50 Hz stimulation (n = 6, p = 0.81, unpaired t-test) and equal initial ATP recovery rates (**j**) after stimulation (p = 0.54, unpaired t-test). **j**, Axonal ATP levels recovered slightly less following 50 Hz stimulation in cKO compared to ctrl (n = 6, p = 0.0498, unpaired t-test). Data are represented as means ± SEM.

**Figure 6.**
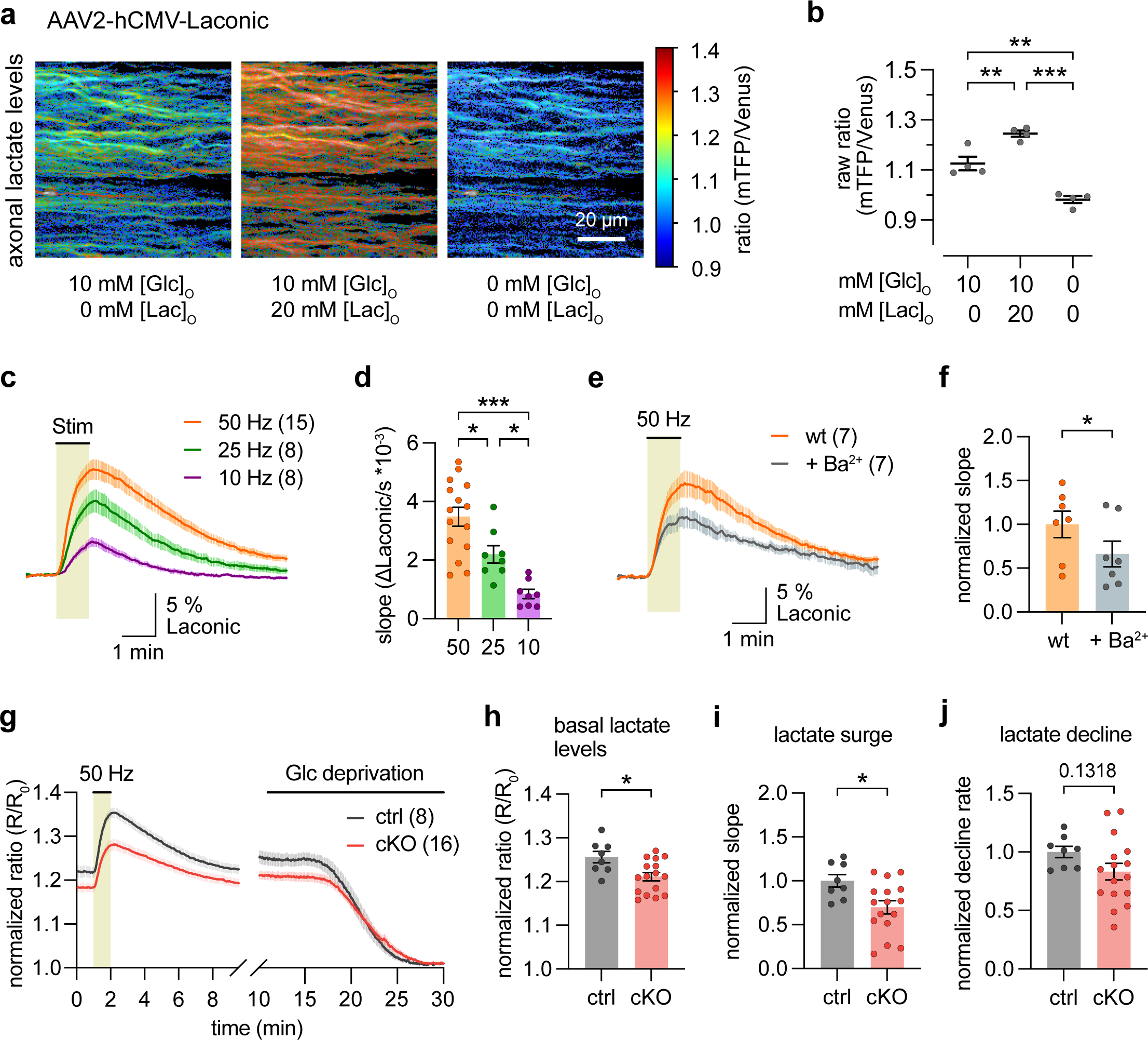
Deficits in axonal lactate dynamics in young Kir4.1 cKO mice. **a,** AAV-mediated lactate FRET sensor (Laconic) expression in optic nerve axons. Depicted are representative color-coded ratio images from a wildtype optic nerve sequentially imaged in regular ACSF with 10 mM glucose (Glc), with additional 20 mM lactate (Lac) and after complete removal of Glc and Lac from the ACSF. Warm and cold colors indicate high and low ratios or lactate levels, respectively. Scalebar 20 µm. **b**, Quantification of ratios obtained from conditions presented in (**a**, in short [Glc]/[Lac] in mM) confirmed that the lactate sensor reliably increased or decreased with the availability of lactate (n = 4 mice, 10/0 vs 10/20: p = 0.0052; 10/0 vs 0/0: p = 0.0014; 10/20 vs 0/0: p < 0.0001; one-way ANOVA Holm-Šídák’s multiple comparisons test). **c**, Axonal lactate levels increase upon high-frequency stimulations. Depicted are lactate level changes (in %) in response to 10 Hz, 25 Hz and 50 Hz stimulations. **d**, Quantification of stimulation-evoked lactate surges (initial slopes) from the recordings in (**c**) showing stronger lactate increases with higher spiking activity (50 Hz vs. 25 Hz, p = 0.0141; 50 Hz vs. 10 Hz, p < 0.0001; 25 Hz vs. 10 Hz, p = 0.0141; one-way ANOVA with Holm-Šídák’s multiple comparisons test). **e** and **f**, Application of Ba^2+^ (100 μM) reduced the stimulation-induced lactate rise in axons of wildtype (wt) mice by 34 ± 11% (n = 7 mice, paired t-test p = 0.0249). **g**, Time course of axonal lactate level changes from optic nerves of ∼3 months old cKO and ctrl mice stimulated with 50 Hz and followed by Glc deprivation (GD). Traces were normalized to the minimum obtained after 20 min of GD. Note that axons from cKO nerves have lower resting lactate levels and stimulus-evoked lactate surges compared to ctrl. **h** and **i**, Basal axonal lactate levels (**h**) (normalized ratios derived from baseline prior to 50 Hz stimulation in **g**) were lower in optic nerves from cKO (n = 16 mice) compared to ctrl (n = 8, p = 0.0119, unpaired t-test), as were 50 Hz-evoked lactate surges (**i**) (p = 0.0174, unpaired t-test). **j**, Lactate decline rates during Glc deprivation (in **g**) were not significantly altered in cKO compared to ctrl (p = 0.1318, unpaired t-test). Data are represented as means ± SEM.

Loss of oligodendroglial Kir4.1 has been reported to cause a decline in axonal integrity with age^23^. At the age of 3 months, visual inspection of electron micrographs did not yield signs of impaired axonal integrity in Kir4.1 cKO mice, and myelin thickness and axonal diameters were not perturbed (**Fig. 3e-g**). However, at the age of 7-8 months we found a significant increase in axon/myelin profiles displaying signs of axonal degeneration (**Fig. 3k and l**) including giant axonal swellings (**Fig. 3m**) in optic nerves from cKO mice. We also found an increase of GFAP immunolabeling in optic nerve sections from 7 months old Kir4.1 cKO mice (**Extended Data Fig. 5c**), consistent with secondary astrocytic reactivity. However, at the same age there was no increase in IBA1 immunopositivity, suggesting absence of overt microgliosis (**Extended Data Fig. 5d**). The age-dependent axonopathy in Kir4.1 cKO mice was not associated with any visible myelin abnormalities or myelin thinning (**Extended Data Fig. 5e and f**), implying that axons may be suffering from a deficit in oligodendroglial support.

Does the absence of oligodendroglial Kir4.1 lead to defective axonal energy metabolism at a young age, before the appearance of axonal pathology? To address this question, we first assessed molecular changes associated with energy metabolism in optic nerves of Kir4.1 cKO mice. We performed tandem mass tag (TMT)-based quantitative proteomics analysis on optic nerve lysates (**Fig. 4a** and **Extended Data Fig. 7a**). As expected, levels of Kir4.1 protein (gene name *Kcnj10*) were significantly reduced in samples from Kir4.1 cKO mice (**Fig. 4b** and **Extended Data Fig. 7b**). Moreover, within the top 50 false discovery rate (FDR)-sorted proteins, we found a reduced abundance in proteins associated with vesicular transport and energy metabolism (**Fig. 4b**). Indeed, overall gene set enrichment analysis (GSEA) and pathway analyses revealed a decrease in transmembrane transporter activity, vesicular transport and oxidative phosphorylation pathways (**Fig. 4c and 4d**).

**Figure 7.**
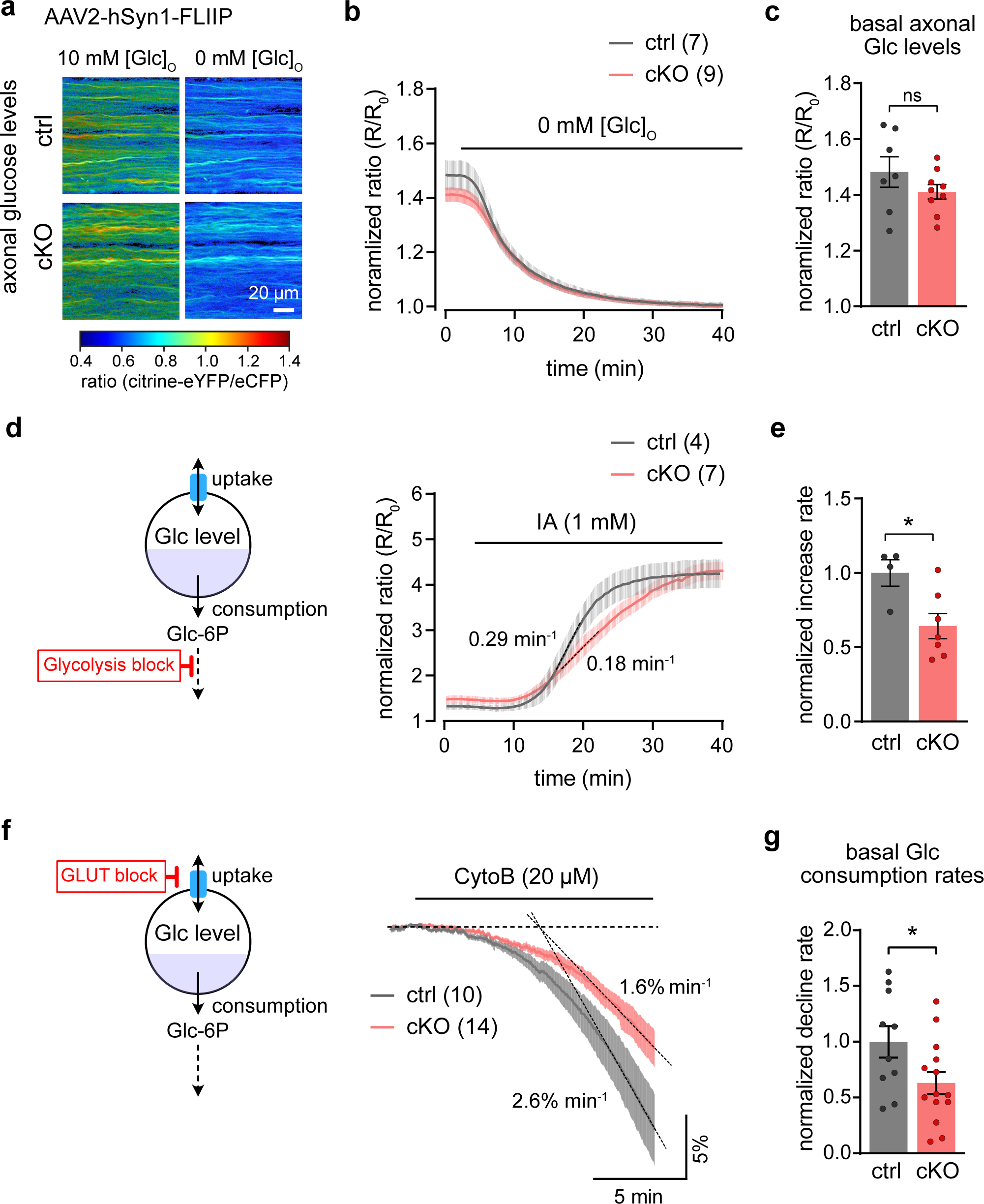
Axonal glucose uptake and metabolism are reduced in young Kir4.1 cKO mice. **a,** AAV-mediated glucose FRET sensor (FLIIP) expression in optic nerve axons. Depicted are representative color-coded ratio images from optic nerves of ctrl (top) and cKO (bottom) in ACSF containing 10 mM glucose (Glc) and after glucose deprivation. Warm and cold colors indicate high and low ratios or glucose levels, respectively. Scalebar 20 µm. **b**, Time course of axonal glucose levels from optic nerves of ∼3 months old cKO and ctrl mice incubated in regular ACSF with 10 mM glucose (Glc) and then switching to zero glucose. Traces were normalized to the minimum obtained after 35 min of glucose deprivation (GD). Of note, 10 mM lactate was included in the ACSF throughout this experiment to sustain axonal conduction during GD and optic nerves were stimulated every 10 s (0.1 Hz) during experiment. **c**, Basal axonal glucose levels (normalized ratios derived from baseline prior to GD in **b**) were comparable between the genotypes (n = 7-9, p = 0.2276, unpaired t-test). **d** and **e**, Inhibition of glycolysis by IA (1 mM) in ACSF containing 10 mM glucose lead to an increase in axonal glucose levels (**d**). Mean increase rates (at dotted lines) are shown for both genotypes. The rate of glucose level increase upon IA treatment (**e**) was reduced in cKO by 36 ± 13 % compared to ctrl (n = 4-7, p = 0.0235, unpaired t-test). **f** and **g**, Assessment of glucose consumption rate by inhibition of glucose uptake using the glucose transporter (GLUT) blocker cytochalasin B (CytoB), while nerves being stimulated every 10 s (0.1 Hz). **f**, Depicted are the time courses of glucose level decline during incubation with 20 µM CytoB for ctrl (n = 10) and cKO (n = 14), with mean decline rates (at dotted lines) of 2.6 ± 0.4%/min and 1.6 ± 0.3%/min, respectively. **g**, Basal axonal glucose consumption rate was reduced in cKO by 37 ± 17% compared to ctrl (p = 0.0384, unpaired t-test). Data are represented as means ± SEM.

Interestingly, among the detected metabolite transporters we found that the abundance of MCT1 (gene name *Slc16a1*; **Fig. 4b** and **Extended Data Fig. 7c**) and GLUT1 (gene name *Slc2a1*; **Extended Data Fig. 7d**) was reduced in optic nerve lysates from cKO mice, but no overt change in glucose transporter GLUT3 (gene name *Slc2a3*; **Extended Data Fig. 7e**). Hence, it appears that loss of oligodendroglial Kir4.1 may have affected the relative abundance of metabolite transporters and energy metabolism in the optic nerve. In fact, MCT1 is expressed by OLs and plays an important role in maintaining long-term axonal integrity^10, 11^, possibly by supplying fast spiking axons with lactate or pyruvate to fuel axonal energy demands^25^. GLUT1 is also expressed by OLs and its abundance in myelin is likely regulated by axonal activity to maintain long-term axonal stability^12^. We therefore tested if the reduced protein levels of MCT1 and GLUT1 in optic nerve lysates was coinciding with a decrease in their abundance in myelin. Indeed, by immunoblotting myelin purified from brains of cKO mice, we found the abundance of both GLUT1 and MCT1 reduced by about half compared to controls (**Fig. 4e**), suggesting that expression and/or surface trafficking of these metabolite transporters in OLs is possibly regulated by K^+^ signaling and Kir4.1 activity.

### Minor changes in axonal ATP dynamics in Kir4.1 cKO mice

We next asked if axonal energy homeostasis was perturbed when oligodendroglial Kir4.1 was lacking, particularly given that metabolite supply to axons could be impaired due to the reduced abundance of GLUT1 and MCT1 and that the GSEA pathway analysis indicated a reduction in pathways related to oxidative phosphorylation. Of note, axonal ATP levels are reduced in a mouse model of spastic paraplegia (*Plp^null/y^* mice) which develop severe axonal pathology^16^. To investigate axonal ATP dynamics in Kir4.1 cKO mice, we expressed the ATP sensor ATeam1.03^47^ via intravitreal AAV-delivery^24^. First, we assessed basal levels of axonal ATP by normalizing the FRET ratios obtained at 0.1 Hz stimulation in ACSF containing 10 mM glucose to the FRET ratios measured after glucose deprivation (GD) and inhibition of mitochondrial respiration (MI) with 5 mM sodium azide (NaN3) to deplete axonal ATP levels (**Figs. 5a-c**). We found no difference in basal axonal ATP levels between the genotypes (**Fig. 5c**). Also, the rate of ATP decline during GD+MI was comparable between the genotypes (**Figs. 5b** and **5d**). Similarly, the rate of ATP recovery following washout of NaN3 and reperfusion with 10 mM glucose was not affected (**Fig. 5e**). Interestingly, we noticed a slightly faster onset of axonal ATP recovery in cKO nerves compared to littermate controls (**Fig. 5b** and **5f**). However, 15-20 min after onset of recovery, axonal ATP levels in control nerves almost completely returned to their initial baseline levels, but axons from cKO nerves recovered less (**Fig. 5b** and **5g**).

The minor deficit in ATP recovery prompted us to further examine the recovery of axonal firing patterns in cKO mice. The block of axonal conduction as seen from the rapid CAP area decline after onset of GD+MI was comparable between the genotypes (**Extended Data Fig. 8a**). Also, the onset and recovery kinetics of axonal firing appeared unchanged (**Extended Data Fig. 8a**). We then evaluated the partial CAP (pCAP) area to determine the dynamics of the first and second CAP peak^24^. The decline rate in pCAP area after the onset of GD+MI was similar between genotypes (**Extended Data Fig. 8b**). However, the recovery dynamics of the pCAP were dissimilar (**Extended Data Fig. 8b**). The pCAP integrates both amplitude and latency of the first two peaks^24^. Closer inspections of the CAP waveforms revealed a striking difference in the recovery kinetics of the CAP peak conduction latency (**Extended Data Fig. 8c**). During recovery in controls, the second CAP peak amplitude slowly increased while the peak latency was decreasing, however, this peak latency shift (increase in conduction velocity) was strongly reduced in cKO nerves (**Extended Data Fig. 8d**). Hence, axons from cKO mice had a deficit in adjusting conduction speed following acute energy deprivation by chemical ischemia. Chemical ischemia is known to elevate extracellular K^+^ concentrations and to alter oligodendroglial K^+^ conductance^48^. It is likely that oligodendroglial K^+^ clearance following the reenergization of axons is involved in quickly adjusting axonal conduction speeds. This deficit in adjusting conduction speed may acutely alter the energy burden on axons, which could probably explain the lower ATP recovery in axons from cKO nerves following chemical ischemia (**Fig. 5b** and **5g**).

**Figure 8.**
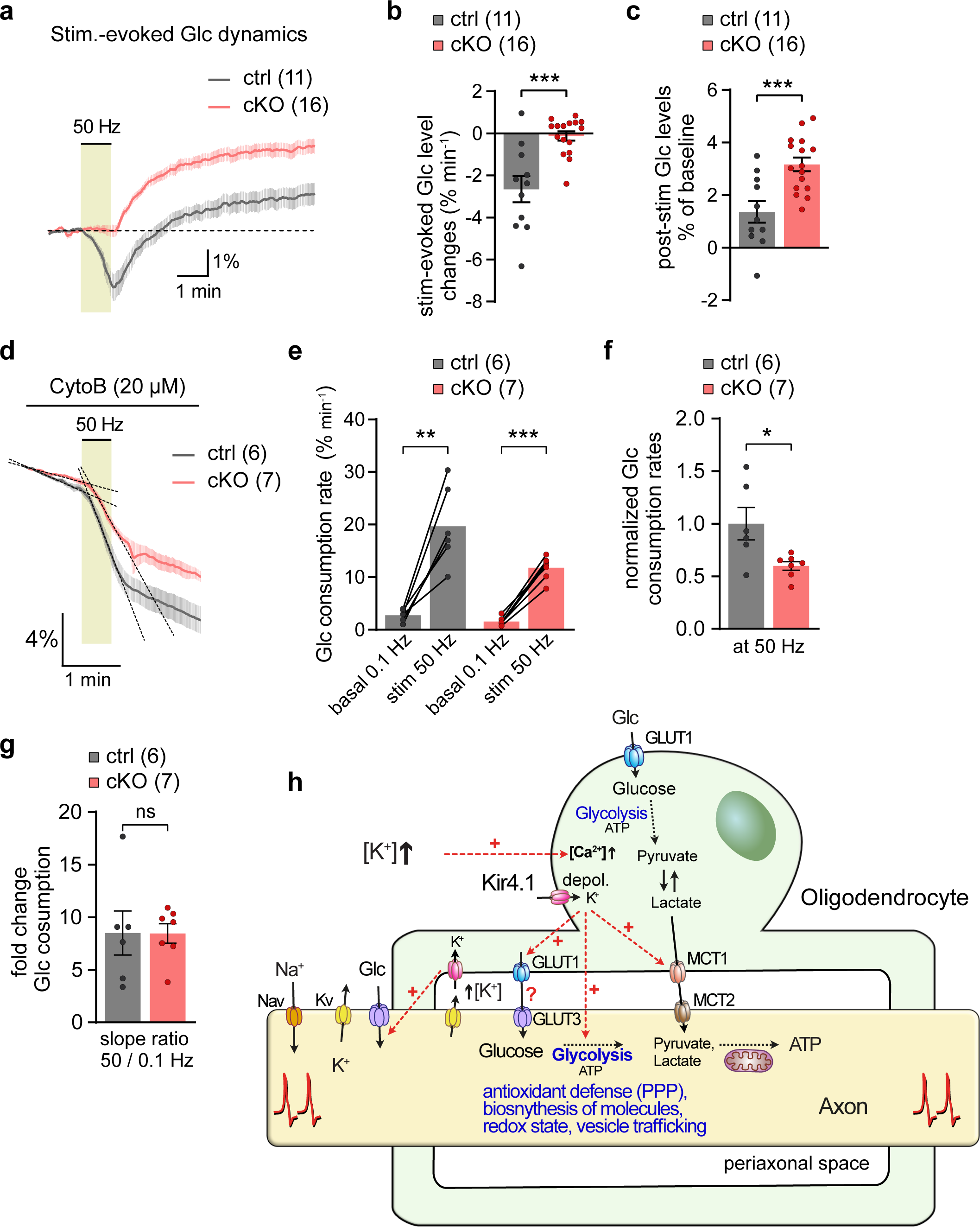
Reduced activity-induced glucose consumption rates in axons from young Kir4.1 cKO mice. **a-c,** Time course traces of 50 Hz stimulation-evoked axonal glucose dynamics (**a**) showing differences in glucose level changes between cKO (n = 16) and ctrl (n = 11). **b**, During stimulation glucose levels decreased at a rate of 2.7 ± 0.6%/min in ctrl but remained stable (0.1 ± 0.2%/min) in cKO (p = 0.0002, unpaired t-test). **c**, After stimulation glucose levels increased above initial baseline values in both genotypes, but significantly higher in cKO (3.2 ± 0.3%) compared to ctrl (1.4 ± 0.4%, p = 0.0006, unpaired t-test). **d-g**, Assessment of glucose consumption rate changes from 0.1 Hz to 50 Hz stimulations in ctrl (n = 6) and cKO (n = 7). Decline slopes are indicated by dashed lines in (**b**). **e**, Axonal glucose consumption rates significantly increased upon 50 Hz in ctrl (0.1 Hz vs 50 Hz, p = 0.0022, paired t-test) and in cKO (0.1 Hz vs 50 Hz, p < 0.0001, paired t-test). **f**, Glucose consumption rate during 50 Hz stimulation was 40 ± 15% lower in cKO compared to ctrl (p = 0.0208, unpaired t-test). **g**, Fold change in glucose consumption from 0.1 Hz to 50 Hz was comparable between the genotypes (8.5 ± 2 in ctrl and 8.5 ± 1 in cKO, p = 0.9858, unpaired t-test). Data are represented as means ± SEM. **h**, Working model in which axon-OL signaling and metabolic coupling in the white matter is controlled by extracellular K^+^ homeostasis via oligodendroglial Kir4.1. Extracellular K^+^ elevations during high frequency axonal firing stimulate a Ca^2+^ rise in OLs that is mediated by membrane depolarization upon K^+^ uptake through Kir4.1 channels. This Kir4.1-mediated axon-OL crosstalk regulates surface expression of MCT1 and GLUT1 in myelin to increase metabolite supply to axons (feed-forward regulation). Besides lactate OLs may also supply axons with glucose, or oligodendroglial Kir4.1 regulates glucose uptake in axons at the nodes of Ranvier. Apart from K^+^ buffering, oligodendroglial Kir4.1 is also in control of axonal glucose metabolism, which is critical for maintaining long-term axonal integrity through various glucose metabolism-dependent processes, including for example antioxidant protection through the pentose phosphate pathway (PPP), biosynthesis of molecules required for structure and function, regulation of the redox state as well as vesicular transport.

Recovery of CAP peak latencies are also reduced following high frequency stimulations in cKO mice (**Extended Data Fig. 6**). We wondered if this would also associate with changes in axonal ATP recovery. We therefore measured ATP dynamics during and after 50 Hz stimulation (**Fig. 5h-k**). Axonal ATP levels dropped during high frequency stimulation with a similar rate and to a similar extent in both cKO and littermate controls (**Fig. 5h** and **5i)**. Also, the initial ATP recovery rate after stimulation was similar between the genotypes (**Fig. 5h** and **5j)**. However, 3-4 min after the onset of recovery, axonal ATP levels recovered slightly less in cKO nerves relative to the initial baseline values (**Fig. 5h** and **5k)**. Hence, in the absence of oligodendroglial Kir4.1, the lower clearance rate of extracellular K^+^ may impose an additional energy cost on axons after high frequency activity or chemical ischemia.

### Early deficits in axonal lactate dynamics in the Kir4.1 cKO mice

Given the reduced abundance of MCT1 and GLUT1 in CNS myelin of Kir4.1 cKO mice, we speculated that axonal lactate fluxes could be affected. Indeed, both of these metabolite transporters were implicated in regulating axonal energy metabolism by supplying axons with lactate or pyruvate^10, 12^. Hence, we next examined if axonal lactate dynamics were altered in Kir4.1 cKO mice. We expressed the lactate sensor *Laconic*^49^ in optic nerve axons following intravitreal AAV-delivery (**Fig. 6a**) and first-tested if the sensor responded to changes in axonal lactate in wildtype nerves. Indeed, raising extracellular lactate levels increased axonal lactate levels (**Fig. 6a** and **6b**) confirming that axons are equipped with transporters for lactate uptake. Removal of glucose and lactate from the ACSF significantly reduced axonal lactate levels (**Fig. 6a** and **6b**). This shows that the *Laconic* sensor in axons is not saturated at baseline (with ACSF containing 10 mM glucose) and its dynamic range would allow the study of lactate dynamics in axons. We then tested if lactate levels change upon increasing axonal firing activity. Indeed, lactate levels increased during high-frequency axonal activity, and the lactate rise was stronger at higher stimulation frequencies (**Fig. 6c** and **6d**). Interestingly, inhibition of Kir4.1 with Ba^2+^ reduced the stimulus-evoked lactate rise in axons by 40% (**Fig. 6e** and **6f**), suggesting that K^+^-mediated axon-OL crosstalk (**Fig. 2**) acutely facilitates lactate supply to axons during spiking activity.

We next examined if axonal lactate dynamics were altered when Kir4.1 was lacking specifically from OLs. Strikingly, both resting lactate levels in axons (at 0.1 Hz stimulation) as well as the high frequency-evoked axonal lactate surge were significantly reduced in Kir4.1 cKO nerves compared to littermate controls (**Fig. 6g-i**). Of note, we compared basal axonal lactate levels between the genotypes after normalizing the FRET ratios to the minimum levels of lactate obtained after 20 min of GD (**Fig. 6g** and **6h**). Lower lactate levels in Kir4.1 cKO axons may result from a reduced lactate supply from OLs, a reduced glycolytic activity in axons and/or an increased axonal lactate consumption rate compared to controls. A higher lactate consumption rate is unlikely given that the decay kinetics of lactate levels during GD were not faster but rather slightly slower in cKO mice compared to littermate controls (**Fig. 6g** and **6j**). This is further corroborated by examining the CAP decline rate during GD, which is identical between the genotypes (**Extended Data Fig. 9a-c**). Moreover, when comparing axonal lactate level changes and CAP performance during GD, both genotypes show the same latency between lactate depletion and CAP decline (**Extended Data Fig. 9d-f**). Notably, it appears that the CAP began to drop only when axonal lactate levels were close to zero (**Extended Data Fig. 9d, e**). Given that the CAP and ATP levels decline almost simultaneously during GD^25^, this strongly suggests that axonal lactate is consumed to maintain axonal ATP levels to fuel action potentials, and once lactate as an axonal energy source is almost depleted, axons stop firing and the CAP starts declining (**Extended Data Fig. 9d, e**). This further implies that axonal mitochondrial respiration is comparable between the genotypes.

### Reduced axonal glucose metabolism in oligodendroglial Kir4.1 cKO mice

The lower lactate levels and activity-mediated lactate surges in axons of Kir4.1 cKO mice reflect a reduced oligodendroglial metabolic support to axons. Both MCT1 and GLUT1 levels were reduced in myelin suggesting that oligodendroglial Kir4.1-mediated K^+^ signaling and clearance are critical in setting-up the metabolic support machinery around axons. In addition to lactate supply, a recent study suggested that callosal OLs may provide axons with glucose^50^. However, little is known about how glucose uptake and glycolysis are regulated in myelinated axons and if OL functions contribute to axonal glucose metabolism. To study glucose dynamics in optic nerve axons we expressed the glucose sensor FLII12Pglu700μΔ6^51^ via intravitreal AAV-delivery (**Fig. 7a**). We first examined basal glucose levels in axons (at 0.1 Hz stimulation) and found comparable levels between the genotypes (**Fig. 7b** and **7c**). Basal glucose levels were obtained by normalizing FRET ratios measured in ACSF containing 10 mM glucose to the minimum glucose levels obtained after 30 min of GD (**Fig. 7b** and **7c**). To make sure that the sensor was not near saturation in axons when bathed with 10 mM glucose, we applied 1 mM iodoacetate (IA) to block glycolysis (**Fig. 7d**). After 35 min of IA treatment, glucose levels increased by 250% and reached the same level in both genotypes (**Fig. 7d**). However, interestingly, we discovered that the rate of axonal glucose rise in cKO nerves was 36% slower than in littermate controls (**Fig. 7d** and **7e**). This may indicate lower glucose uptake rates and/or lower hexokinase activity in cKO axons. To assess glycolytic flux, we blocked glucose transporters with 20 µM Cytochalasin B (CytoB) and measured the slope of cytosolic glucose decline (**Fig. 7f**), as previously described^52, 53^. Strikingly, axonal glucose consumption rate was significantly reduced by 37% in cKO nerves compared to littermate controls (**Fig. 7g**). This suggests that glucose uptake must also be reduced by similar extent in cKO axons to maintain the same basal glucose levels as in controls (**Fig. 7c**), and explains the slower glucose level increase with IA (**Fig. 7d**).

We next-tested if axonal glucose dynamics also differed during high frequency activity. Indeed, when stimulated at 50 Hz, axonal glucose levels decreased in controls but remained constant in cKO (**Fig. 8a** and **8b**). This suggests that activation of glycolysis increases more strongly than glucose uptake in controls but is balanced in cKO nerves. After stimulation, axonal glucose levels increased above baseline in both control and cKO nerves (**Fig. 8a** and **8c**), suggesting that the activation of glucose uptake continued beyond the stimulation period. The difference in glucose dynamics during high frequency activity may result from a lower activation of axonal glucose consumption in cKO compared to controls. To test this, we performed 50 Hz stimulations while blocking glucose import with CytoB. Indeed, high-frequency stimulation increased axonal glucose consumption rates compared to basal (at 0.1 Hz stimulation) glycolytic activity (**Fig. 8d** and **8e**). However, glycolytic activation during 50 Hz stimulation was 40% reduced in cKO compared to littermate controls (**Fig. 8f**). Interestingly, there was an 8-fold increase in the glucose consumption rate upon 50 Hz activity in both control and cKO nerves (**Fig. 8g**). Hence, the machinery for glycolytic activation is not perturbed in axons from Kir4.1 cKO mice, but their overall glucose metabolism is reduced by around 40%, both at rest and during activity. Thus, we conclude that OLs regulate axonal glucose uptake and consumption, a novel metabolic OL-axon interaction that requires Kir4.1 function.

## Discussion

The loss of axonal integrity in white matter in the aging brain and in various neurodegenerative diseases is increasingly considered to result from homeostatic dysfunctions in the axon-OL unit^6, 7, 14, 21, 54–57^. Yet, how OLs interact with and support myelinated axons throughout life is still enigmatic. The results of our study provide a working model in which axon-OL signaling and metabolic coupling in the white matter is controlled by extracellular K^+^ homeostasis, which depends on oligodendroglial Kir4.1 (**Fig. 8h**).

A major correlate of axonal activity, regardless of neuronal subtype, are transient increases in extracellular [K^+^]. OLs are highly permeable to K^+^ and change their membrane potential with changes in external [K^+^]^58^. Indeed, action potentials trigger an inward current and membrane depolarization in OLs, which is facilitated by Ba^2+^-sensitive Kir channels^38, 39^. In this study, we used optic nerve electrophysiology and two-photon Ca^2+^ imaging to study the molecular mechanisms governing white matter axon-OL signaling. Our results suggest that extracellular K^+^ elevations during rapid axonal firing stimulate a Ca^2+^ influx in OL somas, which is mediated by membrane depolarization upon K^+^ uptake through Ba^2+^-sensitive Kir channels. This K^+^-induced depolarization then facilitates Ca^2+^ entry in part through reverse-mode NCX activation. Optic nerve OLs predominantly express Kir4.1 channels^59, 60^, which are particularly sensitive to Ba^2+^. Of note, we found that Ba^2+^ application alone did not change basal Ca^2+^ levels in OLs, however, Ba^2+^ effectively diminished both the stimulus- and K^+^-evoked Ca^2+^ surge specifically in OLs, without affecting axonal Ca^2+^ dynamics. This is consistent with previous electrophysiological recordings indicating that oligodendroglial Kir4.1 is not critical to retain the resting membrane potential of adult white matter OLs, but that Kir4.1 regulates OL membrane depolarization upon axonal activity^22^. Moreover, our CAP recordings from Ba^2+^-treated optic nerves and nerves with OL-specific Kir4.1 deletion revealed equivalent recovery kinetics of axonal conduction after high-frequency stimulation, supporting that Ba^2+^ primarily blocked oligodendroglial Kir4.1-mediated K^+^ uptake. Nonetheless, other Ba^2+^-sensitive K^+^ channels may also be involved in the stimulus- and K^+^-evoked Ca^2+^ response in optic nerve OLs, which requires further investigations in the future. Interestingly, we also observed a consistent undershoot of Ca^2+^ levels in OLs after the stimulations, which was larger with higher stimulation frequency and proportional to the initial rise in Ca^2+^. While the mechanism and function of this post-stimulus Ca^2+^ undershoot needs further exploration, it seems likely that OLs maintain a certain resting Ca^2+^ elevation and that post-stimulus Ca^2+^ removal processes are activated. These may include uptake into mitochondria and endoplasmic reticulum, and activation of plasma membrane Ca^2+^ pumps.

Our results indicate that increases in extracellular K^+^ during high frequency axonal firing activate OL somas in adult white matter. Other activity signals such as glutamate may primarily induce Ca^2+^ activity in specialized microdomains along internodes^61, 62^. Indeed, vesicular glutamate release from axons promote OL precursor cell (OPC) differentiation and to stabilize myelin sheath formation during development^62–67^. Moreover, in response to neuronal activity, myelin sheaths of newly formed OLs in zebrafish exhibit Ca^2+^ signals, which likely regulate myelin sheath growth^68, 69^. In contrast, Ca^2+^ transients in myelin sheaths of developing OLs filled with Oregon Green BAPTA-1 (OGB-1) in mouse cortical slices are independent of neuronal activity^70^. It is possible that axon-OL signaling may vary across developmental stages, anatomical regions or the pattern of neuronal activity. We found that the stimulus-evoked Ca^2+^ response in OLs was proportional to the frequency of axonal spiking and to extracellular K^+^ concentrations. K^+^ is likely released from axons at the nodes of Ranvier^71, 72^ and along the juxtaparanodal domains under the myelin sheath^73, 74^ (**Fig. 8h**). One possibility is that the activity-mediated buildup of extracellular K^+^ around OLs could be more pronounced in fully myelinated, adult white matter tracts compared to more sparsely myelinated cortical regions. This could be resolved in future studies using high-resolution imaging of genetically encoded sensors to determine the spatiotemporal dynamics of activity-induced extracellular K^+^ fluctuations and associated changes in the membrane potential of OLs and their myelin sheaths.

Embryonic deletion of Kir4.1 from both astrocytes and OL lineage cells causes severe white matter pathology, including myelin vacuole formations, loss of myelin integrity, neuroinflammation and axonal loss^22, 46, 75, 76^. These early pathological manifestations are likely explained by the loss of astrocytic Kir4.1 functions, given that OL lineage-specific deletions of Kir4.1 were devoid of myelin abnormalities^22, 23^. We here confirm that oligodendroglial Kir4.1 is not critical for axonal myelination, as myelin sheath thickness and axonal conduction speed were normal in Kir4.1 cKO mice. We also observed no signs of myelin vacuoles, corroborating that K^+^ and water homeostasis is a critical astrocytic function during white matter development^22^. Our findings also revealed that K^+^ clearance via oligodendroglial Kir4.1 is critical for the rapid recovery of axonal firing and conduction speed following high spiking activity, supporting previous observations^22^. In fact, our data suggest that OLs, and not astrocytes, are the primary cells engaged in K^+^ clearance in adult white matter tracts, given that both pharmacological blockage of Kir4.1 and OL-specific Kir4.1 deletion revealed the same CAP recovery dynamics following high-frequency stimulations. On the other hand, astrocytic Kir4.1 is likely critical in controlling extracellular K^+^ dynamics near synapses in close proximity to astrocytic processes^77^. Moreover, OLs and astrocytes are gap-junction coupled and this glial syncytium is essential for K^+^ siphoning during development^39, 73, 75, 78^. However, disruption of the glial network in adult mice did not lead to myelin vacuole formation or axonal pathology^79^. The prevalent notion is that OLs are primarily responsible for K^+^ clearance in adult white matter; however, astrocytes may also be engaged independent of gap junction coupling, presumably by removing K^+^ that passes through OLs to the extracellular space.

By sensing axonal activity through extracellular K^+^ fluctuations, OLs may adjust their metabolic support machinery according to the energetic needs of fast spiking axons. Astrocytes accelerate their GLUT1-mediated glucose uptake^80^ and rate of glucose consumption^53, 81^ in response to transient increases in extracellular K^+^. Moreover, a rise in extracellular K^+^ has been implicated in triggering a rapid lactate release from astrocytes^82, 83^. Conceivably, K^+^ may also stimulate glucose transport, glycolysis and lactate release in white matter OLs. This needs to be resolved in subsequent studies, once AAV-mediated sensor expression in mature OLs is established and feasible in reaching the optic nerve for comparable analysis, also using electrophysiology and two-photon imaging. In the meantime, the observed changes in GLUT1 and MCT1 abundance in myelin of Kir4.1 cKO mice suggest that K^+^ homeostasis is involved in adjusting the gear of the metabolic support machinery. Our previous study revealed that oligodendroglial NMDA receptor signaling stimulates surface expression of GLUT1, suggesting that glutamate activity may adjust the glucose uptake capacity of OLs^12^. Here we show that myelinic GLUT1 and MCT1 are reduced when OLs lack Kir4.1 function. This may indicate that surface trafficking of metabolite transporters in OLs is modulated by K^+^ and Kir4.1-mediated signaling. Interestingly, proteome analysis of optic nerve lysates showed a reduction in proteins involved in intracellular membrane trafficking, including vesicle associated membrane proteins VAMP2 and VAMP3, as well as the small GTPases RAB2B and RAB5A, all of which are expressed by mature OLs^84–86^. While the molecular machinery regulating surface trafficking of metabolite transporters in OLs is still poorly understood, studies on adipocytes showed that VAMP2 and RAB5 family of GTPases are involved in GLUT4 glucose transporter trafficking^87–89^. Of note, VAMP2/3-mediated exocytosis promotes membrane expansion in OLs and myelin formation during development^90^ and VAMP3 regulates surface transport of the myelin protein PLP^91, 92^. In Kir4.1 cKO mice, myelin sheath thickness and abundance of PLP in myelin was not affected, which is not surprising given that other transport-regulating proteins such as VAMP7 are also involved in PLP trafficking^91, 92^. Future research is required to fully understand whether and how K^+^ signaling controls metabolite transporter trafficking in OLs.

Reductions in oligodendroglial GLUT1 or MCT1 were previously associated with late-onset axonal pathology^10–12^. We observed axonopathy in Kir4.1 cKO mice around 7-8 months of age, in line with earlier studies^23^. We therefore reasoned that early deficits in OL metabolite supply to axons may contribute to the age-associated axonal injury in Kir4.1 cKO mice. This was supported by the reduced GLUT1 and MCT1 abundance in myelin of young Kir4.1 cKO mice, about 5 months before the manifestation of axonal damage. Indeed, axonal metabolite imaging revealed reduced basal lactate levels and impaired lactate increases upon high frequency axonal activation. Moreover, by imaging glucose fluxes in axons we uncovered that both glucose uptake and consumption were reduced in young Kir4.1 cKO mice. Hence, the deficits in axonal lactate levels and activity-driven lactate surges likely result from impaired OL lactate supply as well as from deficits in axonal glucose metabolism. Interestingly, we found no changes in resting axonal ATP levels or the speed of ATP replenishment following metabolic challenges, suggesting that the reduction of OL metabolite supply and axonal glucose metabolism did not impact ATP homeostasis. Yet, glucose metabolism is involved in maintaining numerous cell functions beyond ATP production^93, 94^, such as establishing antioxidant protection through the pentose phosphate pathway^95^ or providing glycolytic intermediates for the synthesis of various molecules required for structure and function (**Fig. 8h**). Moreover, axonal glycolysis has been implicated in sustaining fast axonal transport^96^. Hence, deficits in axonal glucose metabolism may impact axonal transport and render myelinated axons more vulnerable to oxidative stress, resulting in axonal damage with age.

Importantly, our study uncovered for the first time that OLs are critically involved in regulating axonal glucose metabolism, i.e. both uptake and consumption. In corpus callosum slice preparations, loading OLs with glucose in conditions of aglycemia could reportedly sustain callosal CAPs, suggesting that OLs are capable of transferring glucose to axons^50^. Given the lower abundance of GLUT1 in myelin from Kir4.1 cKO mice and the accompanying decrease in glucose uptake in axons, it is possible that OLs and myelin constitute a limiting factor for glucose delivery to axons (**Fig. 8h**). Whether OLs regulate glucose supply and/or uptake into axons at the internodal myelin-axon interface or at nodes of Ranvier remains to be assessed. Moreover, how axonal glycolysis is modulated by OL functions is another intriguing question that needs further investigation. OLs could influence axonal energy homeostasis via exosomes^97, 98^, however, it needs to be demonstrated if extracellular K^+^ elevations and Kir4.1 functions are involved in the release of exosomes, similar to glutamate signaling^99^. Alternatively, considering that the Na^+^ pump is implicated in regulating neuronal energy metabolism^100^, changes in the molecular composition of the axonal Na^+^ pumps, possibly due to the impaired K^+^ buffering, may affect glycolysis in axons. Yet, how glucose metabolism of myelinated axons is regulated is still poorly explored and will certainly receive more attention in the future.

In summary, the findings of our study support a model in which axon-OL communication and energy metabolism are controlled by K^+^ signaling and oligodendroglial Kir4.1-mediated regulation of extracellular K^+^ homeostasis.

## Acknowledgments

We thank all lab members for frequent discussions and critical input; Jean-Charles Paterna and the VVF of the Neuroscience Center Zurich (ZNZ) for AAV production; Ari Waisman for MOGiCre mice; the Functional Genomics Center Zurich (FGCZ) for proteomics support; D.E.B and K.-A.N were supported by the Adelson Medical Research Foundation (AMRF). B.W. was supported by the Swiss National Science Foundation (SNFS no. 31003A_156965). A.S.S. was supported by a Synapsis Career Fellowship Award, the Neuroscience Center Zurich (ZNZ), the Cloëtta Foundation, and the Swiss National Science Foundation (SNFS; Eccellenza 187000).

## Author contributions

Conceptualization: Z.J.L. and A.S.S.; Methodology: Z.J.L., L.R., H.B.W., W.M., L.F.B., B.W. and A.S.S.; Investigation: Z.J.L., R.B.J., T.R., and A.S.S.; Resources: D.E.B, L.F.B., K.-A.N., B.W., and A.S.S.; Visualization: Z.J.L., and A.S.S.; Writing: Z.J.L. and A.S.S.; Editing: all authors; Supervision, A.S.S.

## Methods

### Animals

All animal experiments were permitted by the local veterinary authorities in Zurich, in agreement with the guidelines of Swiss Animal Protection Law, Veterinary Office, Canton Zurich (Animal Welfare Act of 16 December 2005 and Animal Welfare Ordinance of 23 April 2008). *PLP-CreERT;RCL-GCaMP6s* mice were generated by crossing PLP-CreERT mice (RRID:IMSR_JAX:005975)^26^ with ROSA26-floxed-STOP-GCaMP6s mice (Ai96; RRID:IMSR_JAX:024106)^28^. Heterozygous *RCL-GCaMP6s* (Ai96) mice were used to express GCaMP6s in optic nerve axons following intravitreal delivery of AAV-Cre. *Kir4.1^fl/fl^;MOGiCre* mice^22^ were obtained from crosses of mice carrying the floxed *Kcnj10* (Kir4.1^fl/fl^)^46^ allele with MOGi-Cre mice^101^. As littermate controls Kir4.1^fl/fl^ mice were used. Transgenic mouse lines were maintained on the C57BL/6 background. For experiments in wildtypes we used Charles River C57BL/6 mice. Both male and female mice were used for experiments. Mice were kept with an inverted 12 hours light/dark cycle and food and water were available ad libitum.

### Tamoxifen treatment

6 to 8 weeks old *PLP-CreERT;RCL-GCaMP6s* mice were injected with tamoxifen to drive GCaMP6s expression specifically in oligodendrocytes. Tamoxifen treatment was done as previously described^79^. Tamoxifen (Sigma, T5648) was dissolved in corn oil (Sigma, C8267) at 10 mg/mL, and was prepared freshly for each experimental cohort. Mice were injected intraperitoneally at a dose of 100 mg/kg body weight per day on three consecutive days. Experiments were conducted starting from around 4 weeks after the first tamoxifen injection.

### Intravitreal AAV injections

Intravitreal injections were carried out according to the procedure described in detail before^24^. In short, mice were anaesthetized with fentanyl (0.05 mg/kg), midazolam (5 mg/kg), and medetomidine (0.5 mg/kg), prepared in NaCl (0.9%) and injected intraperitoneally^102^. Topical application of cyclopentolate (1%) followed by phenylephrine (5%) on each eye was used for pupil dilation. Viscotears liquid gel (transparent) was applied to prevent eyes from drying. Mouse body temperature was kept stable at 37°C using a heating pad. Intravitreal injections were performed under a stereo microscope (SteREO Discovery.V20, Zeiss). An incision was made into the sclera with a 30-gauge (G) needle (insulin syringe, Omnican 50, Braun), a 34G Hamilton syringe filled with AAV suspension was inserted into the opening, and 1.5 μl of virus mix was slowly injected (∼0.1 µl/s) into the vitreous. After intravitreal injection, antibiotic eye drops (ofloxacinum, Floxal, Bausch + Lomb) were applied and the mouse was given the analgesic buprenorphine (0.1 mg/kg BW) or carprofen (10 mg/kg BW). The anesthesia was stopped by intraperitoneal administration of the antagonists [a mixture of atipamezole (2.5 mg/kg) and flumazenil (0.5 mg/kg)]. Mice were monitored closely after the procedure and protected from light as pupils stay dilated for several hours.

### AAV viral vectors

The single-stranded (ss) or self-complementary (sc) AAV vectors used in this study were produced, purified and quantified by the viral vector facility (VVF) of the Neuroscience Center Zurich (ZNZ) as described^103^.

Intravitreal AAV injections were performed with undiluted AAV vectors mixed with fluorescein dye (0.01 mg/ml in PBS). The green fluorescein dye was added to monitor successful injections into the vitreous. The following AAVs and their physical titers, in vector genomes/ml (vg/ml), were used. To induce GCaMP6s expression in optic nerve axons we used *RCL-GCaMP6s* (Ai96) mice injected with scAAV-DJ/2-hCMV-chI-Cre-SV40p(A) [3.4 x 10E12 vg/ml]. To study axonal ATP dynamics we used the FRET sensor ATeam1.03^47^ packaged in ssAAV-2/2-hSyn1-ATeam1.03-WPRE-hGHp(A) [3.0 x 10E12 vg/ml]. Lactate dynamics were studied using the FRET sensor Laconic^49^ packaged in ssAAV-2/2-hCMV-chI-Laconic-WPRE-SV40p(A) [3.0 x 10E12 vg/ml]. Glucose dynamics were studied using the codon-optimized version of the FRET sensor FLII^12^Pglu700μd6^51^, termed here FLIIP, packaged in ssAAV-2/2-hSyn1-FLIIP-WPRE-hGHp(A) [2.9 x10E12 vg/ml].

### Optic nerve electrophysiology and two-photon imaging

We performed acute optic nerve preparations for simultaneous electrophysiology and two-photon imaging as described in detail previously^24^. Briefly, after deep anesthesia with isoflurane and decapitation, optic nerves were removed and gently placed into a custom-modified interface perfusion chamber (Haas Top, Harvard Apparatus). The chamber was perfused with artificial cerebrospinal fluid (ACSF) and maintained at 37°C (TC-10 temperature control system, npi electronic). The solution was constantly bubbled with a mixture of 95% O2 and 5% CO2. The nerve ends were inserted into custom-made suction electrodes filled with ACSF. The optic nerve recording setup was combined with a custom-built two-photon microscope^104^, equipped with a tunable pulsed Ti:Sapphire laser (Chameleon Ultra II; Coherent) and a 25x water immersion objective (XLPLN 25x/1.05 WMP2, Olympus). The microscope was controlled by a customized version of ScanImage (r3.8.1; Janelia Research Campus,^105^). The nerve was allowed to equilibrate in the perfusion chamber with the microscope objective in place for at least 30 min before start of the experiments. A TTL trigger, driven by a stimulus generator (STG4002-1.6mA, Multichannel Systems), was used to synchronize the acquisition of both electrophysiology and imaging.

### Solutions

Optic nerves were superfused with ACSF containing in mM: 126 NaCl, 3 KCl, 2 CaCl2, 1.25 NaH2PO4, 26 NaHCO_3_, 2 MgSO4, 10 glucose, pH 7.4, bubbled with 95% O2 and 5% CO_2_. For pharmacological interventions, drugs at the concentrations mentioned in the text were added to the ACSF shortly prior to experiments. 1000x stock solutions of the following drugs were prepared: TTX (ab120054, Abcam), D-AP5 (#0106, Tocris), PPADS (ab12009, Abcam), Suramin (1472, Tocris), Ouabain (O3125, Sigma-Aldrich), RuR (#1439, Tocris), BaCl_2_ (342920, Sigma-Aldrich), CdCl_2_ (202908, Sigma-Aldrich), NiCl_2_ (339350, Sigma-Aldrich), Iodoacetate (I2512, Sigma-Aldrich), NaN_3_ (S2002, Sigma-Aldrich), NBQX (ab12005, Abcam), 7-CKA (ab120024, Abcam), Nifedipine (#1075, Tocris), Benidipine (#3934, Tocris), Bumetanide (#3108, Tocris), SEA0400 (#6164, Tocris), KB-R7943 (ab120284, Abcam), and Cytochalasin B (#5474, Tocris). When appropriate drugs were protected from light during preparation and the experiment. For analysis of stimulus-evoked Ca^2+^ responses, optic nerves were treated with the drugs for 20-30 min before the experiment. For zero-Ca^2+^ experiments, the ACSF contained 200 μM EGTA and 4 mM Mg^2+^ to maintain constant divalent cation concentration. For [K^+^]_bath_ stimulations, NaCl was adjusted in the ACSF to maintain monovalent cation concentrations. For glucose deprivation experiments, glucose was substituted with sucrose. Similar sodium and osmolarity adjustments were made when sodium L-lactate (Sigma-Aldrich) was added to the ACSF.

### CAP recordings and analysis

Nerves were stimulated with a stimulus generator (STG4002-1.6mA, Multichannel Systems) controlled by MC_Stimulus. Square-wave constant current pulse (50 μs) strength was adjusted to evoke maximum compound action potential (CAP) responses, 0.8 mA were used for all experiments. CAPs were acquired with a USB-ME16-FAI acquisition system (Multichannel Systems) connected to a USB-ME16-FAI preamplifier (gain 4x; Multichannel Systems) and data were collected at 50 kHz using the acquisition software MC_Rack (Multichannel Systems). CAP recordings were analyzed using a custom-written MATLAB script available at GitHub (https://github.com/EIN-lab/CAP-analysis). The stimulus-evoked CAP response usually comprises three peaks, reflecting subgroups of axons with different conduction speeds^12, 24, 106^. During high-frequency stimulations, CAP peak amplitudes normally decrease while peak latencies increase^24^. We analyzed these CAP peak changes focusing the second peak of the CAP. Peak latency was measured as the time between onset of the stimulus artefact and the CAP peaks. Relative changes were calculated by normalizing values to the baseline before high-frequency stimulations. In some experiments, we also measured relative changes in the area under the curve (AUC) of the CAP response, or the partial CAP (pCAP) area which is the AUC of the first and second CAP peaks. The pCAP area integrates changes in amplitude and latency of the first two CAP peaks, likely reflecting changes derived from large and medium-sized axons. The following stimulation protocols were used in this study: Nerves were first stimulated at 0.4 Hz for 1 min to acquire baseline values (e.g. CAP amplitude and Ca^2+^ levels), followed by a 30 s (or 1 min) stimulation period at 10, 25 or 50 Hz and then followed by a recovery period of 4-5 min with 0.4 Hz stimulation. CAP responses were collected every second during high-frequency stimulation periods. In CAP recordings involving glucose deprivation (GI) and inhibition of mitochondrial respiration (MI) nerves were stimulated every 10 s (0.1 Hz) throughout the experiment. We also determined stimulus-response correlations (nerve excitability) by varying the stimulus intensity from 0.1 to 1 mA and measuring the AUC of the corresponding graded CAPs. The CAP area was expressed as percentage of the last CAP area (at maximum stimulation of 1 mA) in the train.

### Calcium imaging and analysis

For GCaMP6s imaging, the sensor was excited at 940 nm with laser powers between 5 and 10 mW. Fluorescence emission was detected with a GaAsP photomultiplier tube (PMT; Hamamatsu Photonics) and a band-pass filter 520/70 nm (Semrock). Overview images were recorded at a resolution of 512 x 512 pixels with a pixel dwell time of 3.2 μs. For Ca^2+^ imaging in OLs or axons, a field of view was selected 15 to 20 μm below nerve surface with 7-8x digital zoom, and images were acquired at 2.96 Hz at a resolution of 128 x 128 pixels with a pixel dwell time of 12.8 μs. Data analysis was carried out with MATLAB (MathWorks, R2015b) using the custom toolbox CHIPS^107^ available on GITHUB (https://github.com/EIN-lab/CHIPS), as previously described^79, 83, 108^. Regions of interest (ROIs) around OL somas were manually defined in ImageJ and ROI masks were fed into the CHIPS analysis pipeline. For axons, the whole frame was used for analysis. Image sequences were motion corrected and a 2D spatial Gaussian filter (σxy = 2 µm) and a temporal moving average filter (width = 1 s) were applied to all images to reduce noise. Background noise was defined as the bottom first percentile pixel value in each frame and was subtracted from every pixel.

The signal vector (dF/F) from each ROI (or whole frame) was calculated using the mean intensity of 10 frames before the onset of the stimulation paradigms as baseline. The evoked Ca^2+^ responses were quantified as the AUC of the signal change during the 30 s stimulation period. Pharmacological effects were evaluated for each individual cell stimulated before and after drug application, and individual responses were averaged from 2-3 imaging sessions. This paired-analysis approach allowed to reduce variability between cells from different recordings and nerves. The number of mice (N) and cells (n) are indicated in the figure legends.

### Metabolite imaging and analysis

The FRET-based metabolite sensors (Laconic, ATeam1.03 or FLIIP) were excited at 870 nm with laser powers between 5 and 15 mW, and donor and acceptor fluorescence signals were collected simultaneously with two PMTs using a dichroic beam-splitter (560 nm edge, BrightLine; Semrock) and two band-pass filters 545/55 nm (yellow channel) and 475/50 nm (blue channel). Images were acquired every 2 s or 10s at a resolution of 256 x 256 pixels and a pixel dwell time of 6.4 μs. FRET analysis was performed with a custom-written MATLAB script. The acquired image time-series was time-smoothed using a moving average over 5 images. The number of images for smoothing was carefully selected after testing various time windows that did not affect the temporal dynamics of the acquired image time-series. Background areas were removed by thresholding and the average of the whole frame intensity was extracted for each channel. The ratio of donor to acceptor (Laconic) or acceptor to donor (ATeam1.03 and FLIIP) was calculated and normalized to the corresponding baseline or quasi zero time point as stated in the text. To visualize axonal structures while preserving the quantitative information about the FRET ratio, we used the following strategy: a color scale was applied to the ratio images calculated as pixel-by-pixel division of the two channels after averaging of 20 frames. The generated RGB images were then converted in the YCbCr color space, in which the Cb and Cr coordinates encode for the color information, while the Y one encodes the brightness. The Y coordinate was then replaced with the square root of the sum of the donor and acceptor images, and the whole YCbCr image was converted back to RGB.

### Immunohistochemistry

Mice were anesthetized with pentobarbital and perfused with 2% paraformaldehyde (PFA, Paraformaldehyde Granular, Electron Microscopy Sciences, Hatfield, PA) in PBS. Post-fixation of the optic nerves was done in 4% PFA (in PBS) for 1 hour. Nerves were embedded in frozen section medium (Richard-Allan Scientific NEG 50, Thermo Scientific), cut longitudinally in sections of 12 μm with a cryostat (CM3050 S, Leica) and transferred onto SuperFrost Plus slides (Thermo Scientific). Immunohistochemical labeling was performed directly on the slides. Sections were treated with 0.3% Triton X-100 (Sigma-Aldrich, Buchs, Switzerland) in Tris buffer (50 mM, pH 7.4) containing 4% normal donkey serum for 1 hour at room temperature (RT). Primary antibodies (see Table 1) were incubated in same solution over night at 4 °C. Secondary antibodies raised in donkey were diluted 1:700 in 0.05% Tris-Triton containing 4% normal donkey serum and applied for one hour at RT. DAPI (1:10000 from 1 mg/ml stock, Thermo Fisher Cat. D3571), used for nuclei labeling, was added to the secondary antibodies. Confocal images and z-stacks of 10 µm were acquired with a Zeiss LSM 700 or Zeiss LSM 800 confocal laser scanning microscope equipped with a 40× objective (Plan-Apochromat, NA 1.4, Oil DIC (UV) VIS-IR). Image analysis was performed with ImageJ (Fiji version 1.52p). For GFAP and IBA1 analysis, maximum intensity projections were binarized and the fluorescent particle area was determined. For quantifications, 2 images per section and 4 sections per animal were analyzed.

**Table 1.** Antibody information. IHC, immunohistochemistry, IB, immunoblot

| Antigen | Host Species | Method, dilution | Source and Cat# |
| --- | --- | --- | --- |
| α-CC1 | mouse, monoclonal | IHC, 1:100 | Calbiochem, Cat# OP80 |
| α-GFP | chicken, polyclonal | IHC, 1:1000 | Aves Labs, Cat# GFP-1020 |
| α-GFAP | chicken, polyclonal | IHC, 1:2000 | Abcam, Cat# ab4674 |
| α-IBA1 | rabbit, polyclonal | IHC, 1:1000 | FUJIFILM Wako Chemicals, Cat# 019-19741 |
| α-Kir4.1 | rabbit, polyclonal | IB, 1:1000 | Alomone, Cat# APC-035 |
| α-MCT1 / SLC16A1 | rabbit, polyclonal | IB, 1:500 | prod. by Kathrin Kusch<sup>115</sup> |
| α-GLUT1 | rabbit, polyclonal | IB, 1:500 | prod. by Kathrin Kusch<sup>116</sup> |
| α-CNP | mouse, monoclonal | IB, 1:1000 | Sigma, Cat# C 5922 |
| α-PLP | rabbit, polyclonal | IB, 1:5000 | A431<sup>117</sup> |
| α-MOG (8-18C5) | mouse, monoclonal | IB, 1:5000 | Christopher Linnington |
| α-ATP2a1 | mouse, monoclonal | IB, 1:1000 | Abcam, Cat# ab7671 |
| α-ATP1a3 | mouse, monoclonal | IB, 1:1000 | Abcam, Cat# ab2826 |
| α-mouse IgG HRP | goat, polyclonal | IB, 1:10000 | Jackson ImmunoResearch, Cat# 115-035-003 |
| α-rabbit IgG HRP | goat, polyclonal | IB, 1:10000 | Jackson ImmunoResearch, Cat# 111-035-003 |
| α-mouse Cy3 | donkey | IHC, 1:700 | Lucerna Chem, Cat# 715-165-151 |
| α-rabbit Cy3 | donkey | IHC, 1:700 | Jackson ImmunoResearch, Cat# 711-165-152 |
| α-chicken Alexa 488 | donkey | IHC, 1:700 | Jackson ImmunoResearch, Cat# 711-545-152 |

### Electron microscopy and analysis

Mice were deeply anaesthetized using isoflurane before decapitation and optic nerve extraction. Optic nerves were immediately transferred to fixative solution (4% PFA, 2.5% glutaraldehyde in phosphate buffer with 0.5% NaCl, pH 7.4) and fixed overnight at 4°C. Tissue preparation and electron microscopy (EM) were carried out as previously described^109^. In short, following post-fixation with 2% OsO4 (Science Services, Munich, Germany) in 0.1 M phosphate buffer pH 7.3 and acetone dehydration, tissue fragments were embedded in EPON (Serva, Heidelberg, Germany). Ultrathin sections were cut with a Leica UC7 ultramicrotome (Leica, Vienna, Austria) and then stained using UranylLess™ (Science Services, Munich, Germany). EM pictures were captured with a Zeiss EM912 electron microscope (Carl Zeiss Microscopy GmbH, Oberkochen, Germany) equipped with an on-axis 2k CCD camera (TRS, Moorenweis, Germany). EM image analysis was performed using ImageJ (Fiji, version 1.52p). For assessment of axonal diameters and g-ratios (ratio of axon diameter to axon diameter including the myelin sheath), 5 random overview EM pictures (at 8000× magnification) from optic nerve sections were taken and a total of 250 axons per animal were analyzed. Axonal diameter and g-ratios were calculated from circular areas equivalent to the measured areas. For assessment of axonal pathology, 8-10 EM images were analyzed per animal. The experimenters were blinded to the genotypes.

### Proteomics and analysis

Optic nerves from 2.5 months old Kir4.1 cKO and littermate controls were extracted following deep anesthesia using isoflurane. Tissue was homogenized in PreOMICS lysis buffer using 0.5mm Zirconium Oxide beads (ZROB05, Next Advance, Troy, NY) and the tissue homogenizer Bullet Blender (BBX24, Next Advance; 2 x 15 s cycles on setting 10). The TMT-based quantitative proteomics was conducted by the Functional Genomics Center Zurich (FGCZ) as previously described^79^. Briefly, protein concentrations were determined using the Lunatic UV/Vis polychromatic spectrophotometer (Unchained Labs). 100 μg of protein from each sample were reduced with 2 mM TCEP (tris(2-carboxyethyl)phosphine) and alkylated with 15 mM iodoacetamide at 60°C for 30 min. Samples were processed using the single-pot solid-phase enhanced sample preparation. For TMT-labelling, 200 μg TMT10plex reagent (Thermo Fisher Scientific, 90110) were mixed with 41 μl of anhydrous acetonitrile (Sigma-Aldrich) and added to 45 μg peptides in 100 μl of 50 mM TEAB, pH 8.5. After the solution was mixed and incubated for 60 min at room temperature, 8 μl of 5% hydroxylamine (Thermo Fisher Scientific) was added to quench the reaction. To get the combined TMT sample, equal amounts of each TMT channel were mixed. Using high pH reveres phase chromatography, labeled peptides were offline pre-fractionated. A new fraction was obtained every minute and concatenated to 8 final fractions. These were then dried down and resuspended in 40 μL of 3% acetonitrile, 0.1% formic acid containing indexed retention time (iRT)-peptides (Biognosys). For mass spectrometry analysis, an Orbitrap Fusion Lumos (Thermo Scientific) was used, which was equipped with a Digital PicoView source (New Objective) and coupled to a M-Class UPLC (Waters). Channel A had a solvent composition of 0.1% formic acid and channel B of 0.1% formic acid and 99.9% acetonirile. 600 ng of peptides were loaded per sample on a commercial MZ Symmetry C18 Trap Column (100 Å, 5 μm, 180 μm × 20 mm, Waters) followed by nanoEase MZ C18 HSS T3 Column (100 Å, 1.8 μm, 75 μm × 250 mm, Waters). A quadrupole isolation with a window of 0.7 Da and HCD fragmentation with 38% fragmentation energy was used to record data-dependent MS/MS in the Orbitrap. A minimum intensity threshold of 50’000 was used for MS/MS and the cycle time was set to a maximum of 3 s. The local laboratory information management system (LIMS;^110^) was used to handle the mass spectrometry proteomics data. Proteome Discoverer (PD version 2.4) was used to analyze the raw MS data, before protein identification with the integrated Sequest HT search engine. Spectra were compared to the mus musculus reference proteome (downloaded from UniProt, 20190709), concatenated with common protein contaminants. Enzyme specificity was set to trypsin tolerating a minimal peptide length of 6 amino acids and a maximum of two missed-cleavages. The default tolerance parameters of PD were used. A maximal false discovery rate (FDR) of 0.01 was set for peptides. For reporter ion quantification the integration tolerance for the most prominent peak was 20 ppm. The intensity values listed in the Protein output were used to compute protein fold changes. To find proteins having two or more peptides, a set of functions implemented in the R package prolfqua^111^ was employed. A modified robust z-score transformation that keeps the original data variance was utilized to normalize the data. To find differences between the genotypes a linear model was fitted to all proteins, contrasts were calculated and the variance, t-statistics, and p-values were moderated^112^. The Benjamini-Hochberg adjustment was used to calculate FDRs from p-values. Furthermore, gene set enrichment analysis (GSEA) was carried out with the WEB-based GEne SeT AnaLysis Toolkit (WebGestalt.org)^113^.

### Myelin purification and immunoblotting

Myelin-enriched light-weight membrane fraction was purified from brains of 2.5 months old Kir4.1 cKO and littermate controls, using sucrose density centrifugation and osmotic shocks as previously described^114^. Brain tissue was extracted after decapitation following deep anesthesia using isoflurane. Protein concentrations of brain lysate and myelin fractions were determined using the DC Protein Assay Kit (Bio-Rad, Munich, Germany) following the manufacturer’s instruction and measured using the Eon High Performance Microplate Spectrophotometer (BioTek, Vermont, USA). Immunoblotting was carried out as previously outlined^18^. Myelin fraction samples were diluted in 4x sodium dodecyl sulfate (SDS) sample buffer (glycerol 40% [w/v], Tris/HCl pH 6.8 240 mM, SDS 8% [w/v] bromophenol blue 0.04% [w/v]); with 5% dithiothreitol (DTT) was added as a reducing agent. Before usage, samples were heated at 40°C for 10 min. For protein separation by SDS-PAGE, the Mini-PROTEAN Handcast system (Bio-Rad, Munich, Germany) was used with self-casted acrylamide gels (10–15%). 5–15 µg samples were loaded per well (depending on protein of interest) next to 5 µl pre-stained protein ladder (PageRuler, Thermo Fisher Scientific, Waltham, USA). Proteins were separated by constant current (200 V) for 45–60 min using a Bio-Rad power supply. Immunoblotting was carried out with a Novex Semi-Dry Blotter (Invitrogen, Karlsruhe, Germany) and proteins were transferred to an activated (100% ethanol, 1 min; followed by two washing steps with water) PVDF membrane (Roche Diagnostics GmbH, Mannheim; Cat# 03010040001) at 20 V for 40 min. After blotting, membranes were blocked in 1x TBS containing 5% non-fat dry milk (Frema, Karlsruhe, Germany) and 0.05% Tween-20 for 1 hour at room temperature. Primary antibodies were diluted in 5 ml blocking buffer and incubated overnight at 4°C and horizontal rotation. Membranes were washed thrice with TBS-T for 5–10 min each and incubated for 1 hour with secondary HRP antibodies diluted in blocking buffer. Membranes were washed three times with TBS-T for 5–10 min. Detection was carried out using enhanced chemiluminescent detection (ECL) according to the manufacturer’s instructions (Western Lightning Plus-ECL or SUperSignal West Femto Maximum Sensitive Substrate; Thermo Fisher Scientific, St Leon-Rot, Germany). Immunoblots were scanned using ECL Chemostar (Intas Science Imaging, Göttingen, Germany). For antibody information, see Table 1.

### Statistical analyses

Inter-group comparisons were made by paired t-test, Mann Whitney t-test or two-tailed Student’s t-test as stated in the figure legends. For multiple comparisons, data were analyzed with one-way or two-way ANOVA with Bonferroni’s post-test. All analyses were conducted through GraphPad Prism 9 or R (v.3.2.2, R Core Team, 2015). The levels of significance were set as *p < 0.05; **p < 0.01; ***p < 0.001. All data is presented as mean ± SEM. P values together with sample size are stated in the figure legends.

---

**Extended Data Figure 1.**
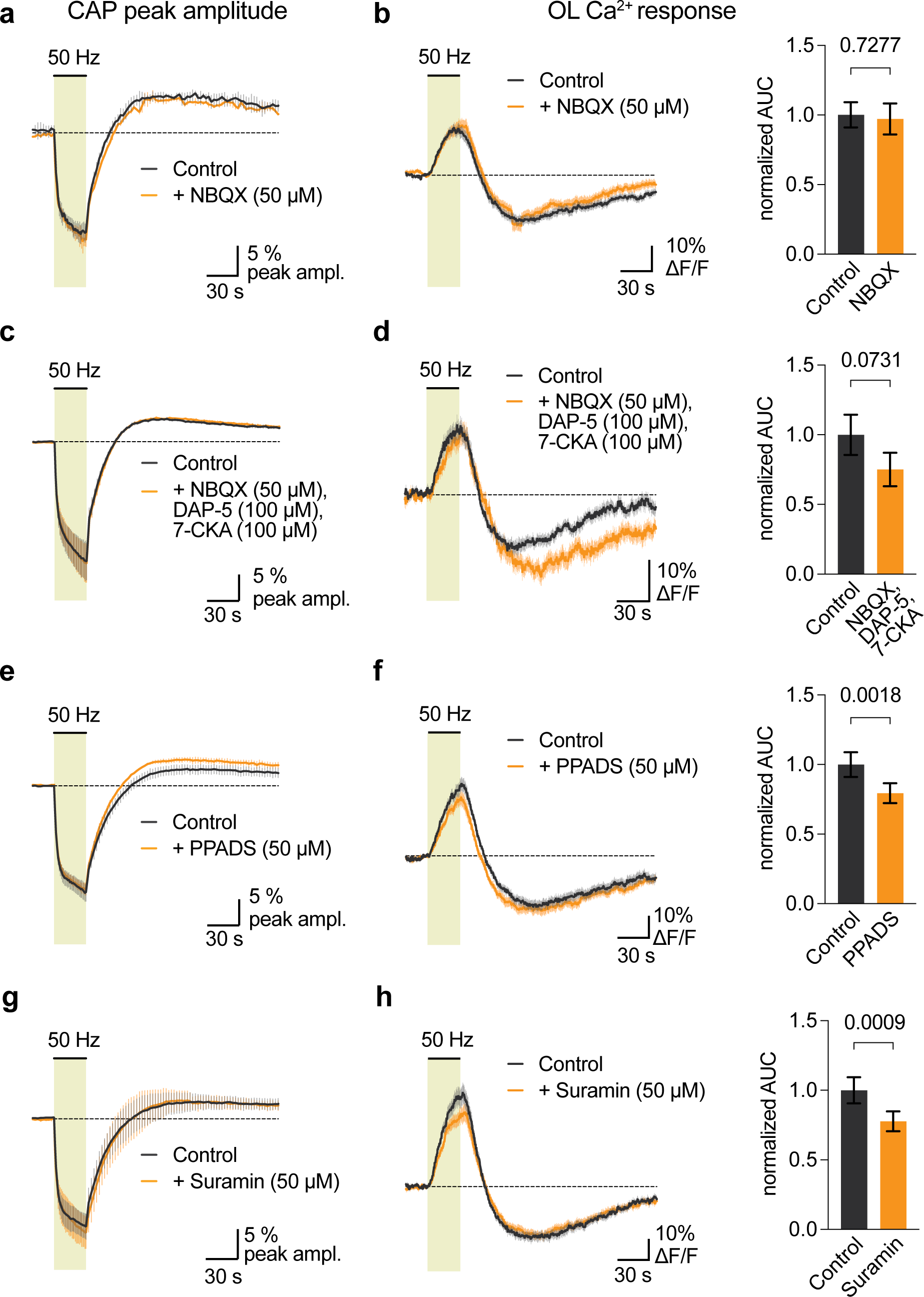
Minor contribution of glutamatergic and purinergic signaling to stimulus-evoked Ca^2+^ increases in OLs. **a** and **b,** Depicted are (**a**) CAP peak amplitude changes and (**b**) OL soma Ca^2+^ changes evoked by 50 Hz stimulations in control condition and in the presence of NBQX (50 µM). Quantification of the OL Ca^2+^ surge (as AUC during stimulation period, bar graphs) revealed no difference in the stimulus-evoked Ca^2+^ increase (n = 45 cells, N = 3 optic nerves, p = 0.7277, paired t-test). **c** and **d,** Analysis of (**c**) CAP peak amplitude changes and (**d**) OL soma Ca^2+^ changes evoked by 50 Hz stimulations in control condition and in the presence of NBQX (50 µM) DAP-5 (100 µM) and 7-CKA (100 µM), revealed a decrease in the stimulus-evoked Ca^2+^ increase by 25 ± 13% (n = 33 cells, N = 3 optic nerves, p = 0.0732, paired t-test). **e** and **f,** Analysis of (**e**) CAP peak amplitude changes and (**f**) OL soma Ca^2+^ changes evoked by 50 Hz stimulations in control condition and in the presence of PPADS (50 µM) revealed a decrease in the stimulus-evoked Ca^2+^ increase by 21 ± 6% (n = 46 cells, N = 3 optic nerves, p = 0.0018, paired t-test). **g** and **h,** Analysis of (**g**) CAP peak amplitude changes and (**h**) OL soma Ca^2+^ changes evoked by 50 Hz stimulations in control condition and in the presence of Suramin (50 µM) revealed a decrease in the stimulus-evoked Ca^2+^ increase by 22 ± 6% (n = 33 cells, N = 2 optic nerves, p = 0.0009, paired t-test). Data are represented as means ± SEM.

**Extended Data Figure 2.**
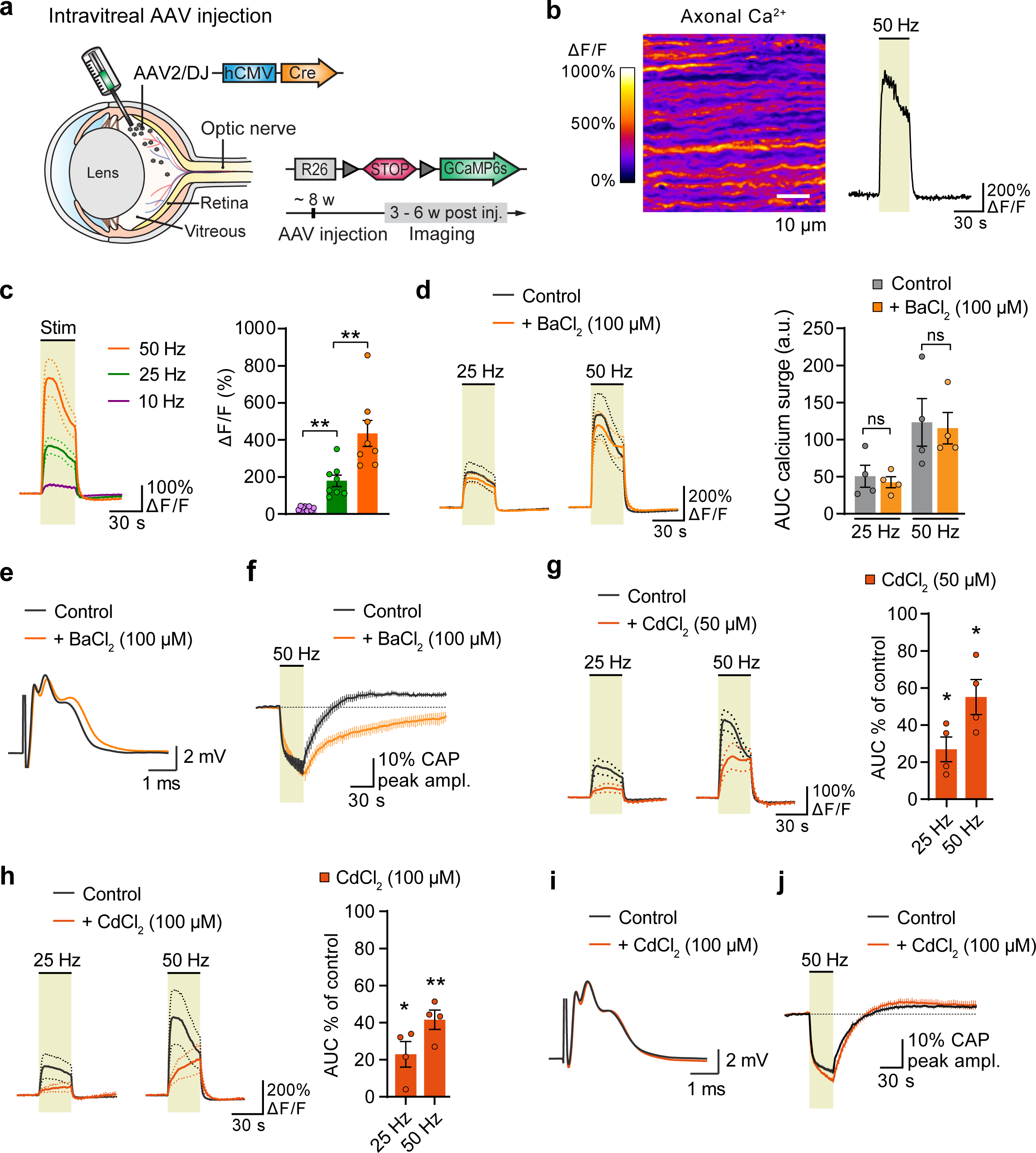
Stimulus-evoked axonal Ca^2+^ surges are not sensitive to barium. **a,** Intravitreal AAV-Cre injection into *RCL-GCaMP6s* mice at the age of around 8 weeks drives abundant GCaMP6s expression in optic nerve axons 3-6 weeks later, allowing for two-photon imaging experiments. **b**, Electrical 50 Hz stimulation triggers a strong Ca^2+^ surges in optic nerve axons. Depicted is an example recording of a 30 s stimulation, with ΔF/F image (left) and corresponding Ca^2+^ trace. **c**, Stimulus-evoked Ca^2+^ responses at different stimulation frequencies. Evoked Ca^2+^ transients are significantly larger with higher stimulation frequencies (10 Hz vs 25 Hz, p = 0.0027; 25 Hz vs 50 Hz, p = 0.0013; one-way ANOVA with Tukey’s multiple comparisons test). **d**, The stimulus-evoked Ca^2+^ surge in optic nerve axons is not affected by application of 100 µM Ba^2+^, tested at 25 Hz (n = 4, p = 0.49, paired t-test) and 50 Hz (n = 4, p = 0.6859, paired t-test) stimulation frequencies. **e**, Example CAP trace in control conditions and with addition of 100 µM Ba^2+^. **f**, Time course of CAP peak amplitude changes upon 50 Hz stimulations. Note that the recovery kinetics of the peak amplitude is strongly reduced in the presence of Ba^2+^ compared to control (n = 4, *F_interaction_* (74, 444) = 2.487, p < 0.0001, two-way ANOVA) **g** and **h**, Inhibition of VGCCs with Cd^2+^ significantly reduced the stimulus-evoked axonal Ca^2+^ surge. **g**, At 50 µM Cd^2+^ the 25 Hz- and 50 Hz-evoked Ca^2+^ surges were reduced to 27 ± 7% (p = 0.0133, paired t-test) and to 55 ± 9% (p = 0.0343, paired t-test), respectively. **h**, At 100 µM Cd^2+^ the 25 Hz- and 50 Hz-evoked Ca^2+^ surges were reduced to 23 ± 7% (p = 0.0402, paired t-test) and by 42 ± 5% (p = 0.0074, paired t-test), respectively. **i,** Example CAP trace in control conditions and with addition of 50 µM Cd^2+^. **j**, CAP recovery kinetics after 50 Hz stimulation is not affected by Cd^2+^. Data are represented as means ± SEM.

**Extended Data Figure 3.**
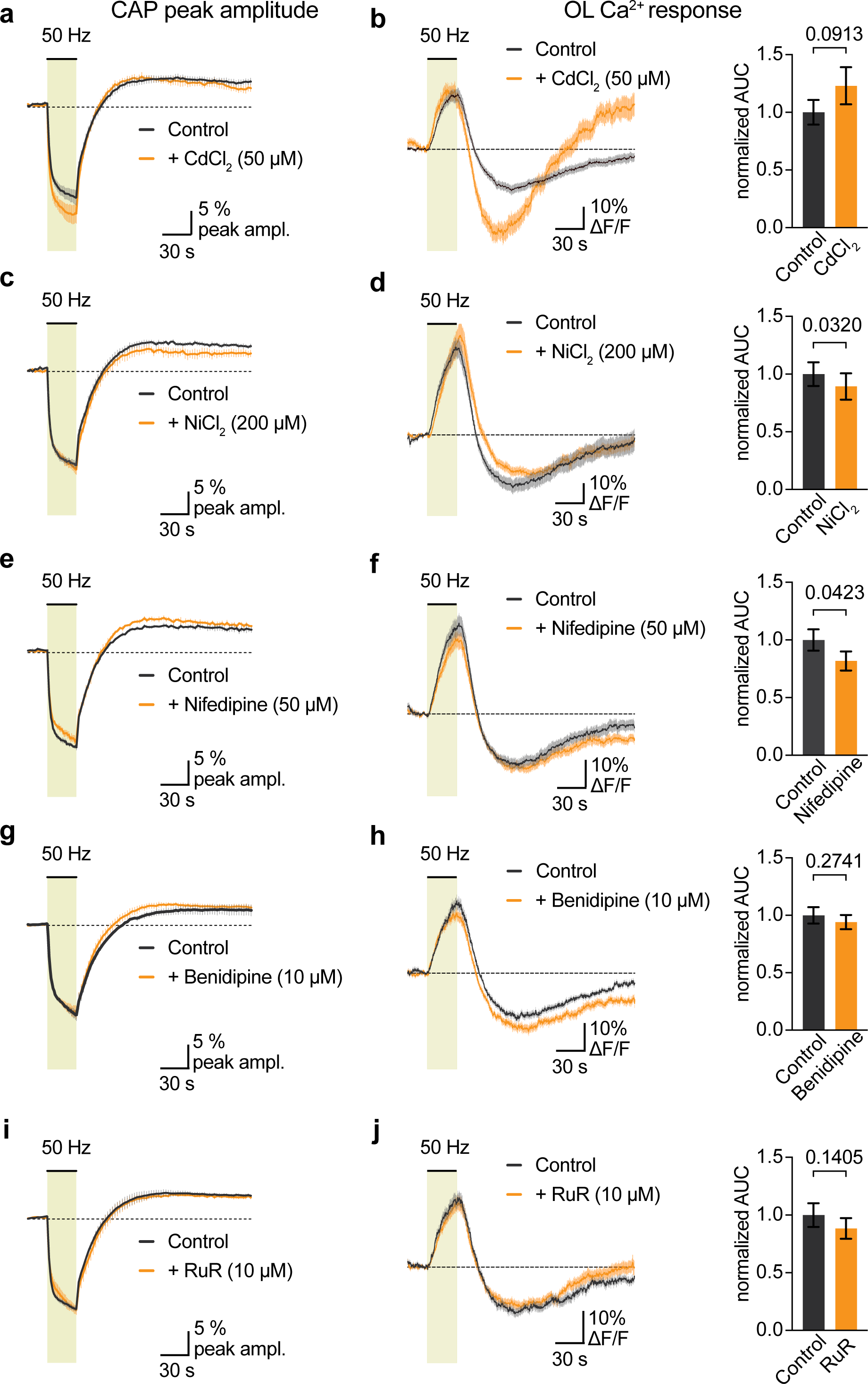
Voltage-gated Ca^2+^ channels are no major drivers of stimulus-evoked OL Ca^2+^ response. **a** and **b,** Analysis of (**a**) CAP peak amplitude changes and (**b**) OL soma Ca^2+^ changes evoked by 50 Hz stimulations in control condition and in the presence of Cd^2+^ (50 µM), revealed a slight increase in the stimulus-evoked Ca^2+^ rise (n = 54 cells, N = 4 optic nerves, p = 0.0913 paired t-test). **c** and **d,** Analysis of (**c**) CAP peak amplitude changes and (**d**) OL soma Ca^2+^ changes evoked by 50 Hz stimulations in control condition and in the presence of Ni^2+^ (200 µM), revealed a slight decrease in the stimulus-evoked Ca^2+^ surge by 11 ± 5% (n = 60 cells, N = 4 optic nerves, p = 0.0320 paired t-test). **e** and **f,** Analysis of (**e**) CAP peak amplitude changes and (**f**) OL soma Ca^2+^ changes evoked by 50 Hz stimulations in control condition and in the presence of Nifedipine (50 µM), revealed a slight decrease in the stimulus-evoked Ca^2+^ surge by 18 ± 9% (n = 39 cells, N = 3 optic nerves, p = 0.0423 paired t-test). **g** and **h,** Analysis of (**e**) CAP peak amplitude changes and (**f**) OL soma Ca^2+^ changes evoked by 50 Hz stimulations in control condition and in the presence of Benidipine (10 µM), revealed no overt change in the stimulus-evoked Ca^2+^ surge (n = 56 cells, N = 3 optic nerves, p = 0.2741 paired t-test). **i** and **j,** Analysis of (**e**) CAP peak amplitude changes and (**f**) OL soma Ca^2+^ changes evoked by 50 Hz stimulations in control condition and in the presence of RuR (10 µM), revealed no significant changes in the stimulus-evoked Ca^2+^ surge (n = 71 cells, N = 4 optic nerves, p = 0.1405 paired t-test). Data are represented as means ± SEM.

**Extended Data Figure 4.**
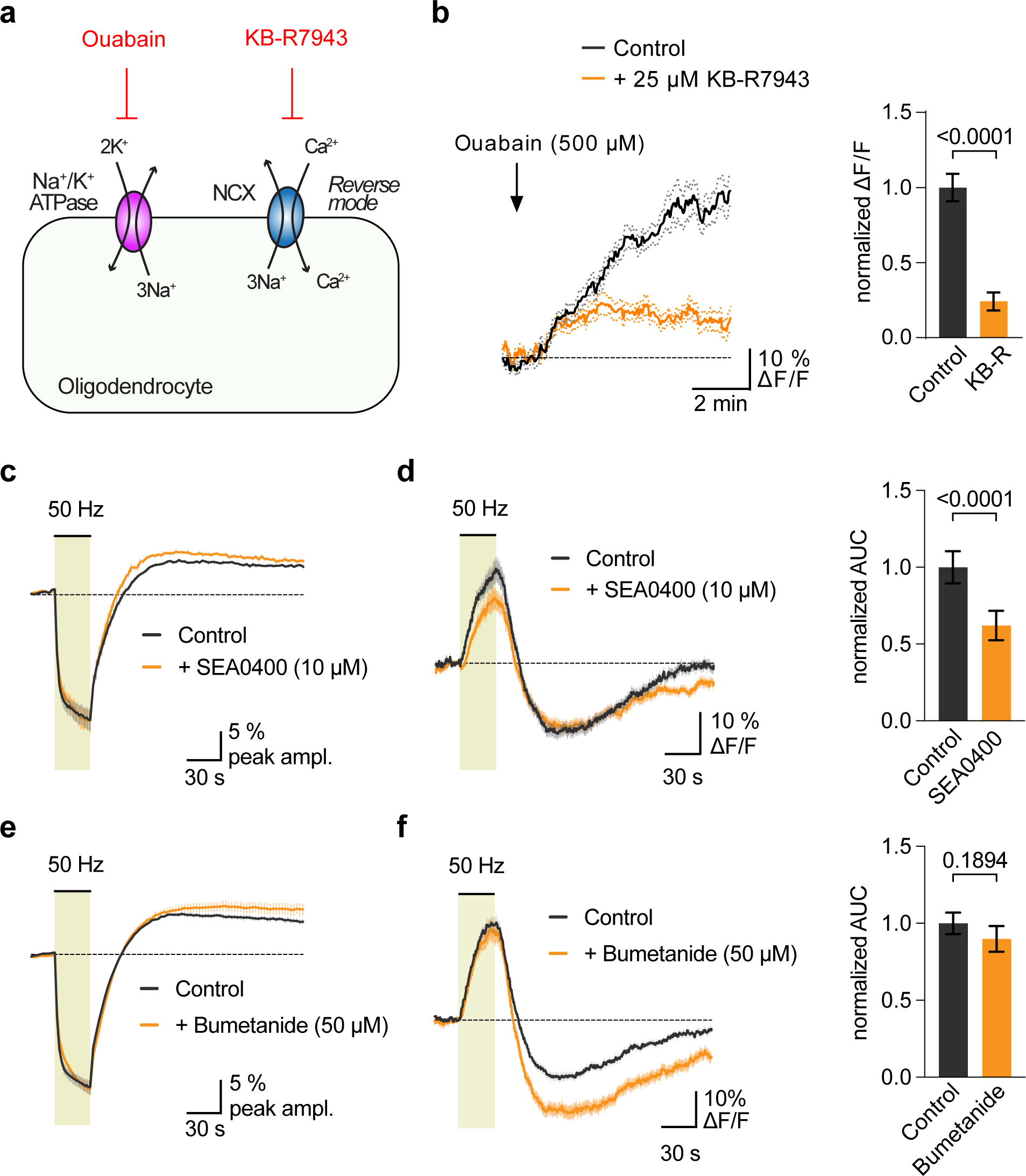
Reverse-mode NCX activation in OLs contributes to the stimulus-evoked OL Ca^2+^ response. **a,** Scheme of testing reverse-mode NCX activity in OLs. Blocking the sodium pump with Ouabain raises intracellular Na^+^ concentration which should activate NCX to drive Na^+^ out and Ca^2+^ in. And by blocking NCX with e.g. KB-R7943, this oubain-evoked Ca^2+^ rise should be reduced. **b**, Indeed, 500 µM oubain application increased Ca^2+^ levels in OLs that could be strongly reduced by addition of 25 µM KB-R7943 (n = 31 cells, N = 2 optic nerves, p < 0.0001, t-test). **c** and **d,** Analysis of (**c**) CAP peak amplitude changes and (**d**) OL soma Ca^2+^ changes evoked by 50 Hz stimulations in control condition and in the presence of SEA0400 (10 µM), revealed a significant decrease in the stimulus-evoked Ca^2+^ surge by 38 ± 7% (n = 54 cells, N = 4 optic nerves, p < 0.0001, paired t-test). **e** and **f,** Analysis of (**e**) CAP peak amplitude changes and (**f**) OL soma Ca^2+^ changes evoked by 50 Hz stimulations in control condition and in the presence of Bumetanide (50 µM), revealed no overt change the stimulus-evoked Ca^2+^ surge by (n = 77 cells, N = 4 optic nerves, p = 0.1894, paired t-test). Data are represented as means ± SEM.

**Extended Data Figure 5.**
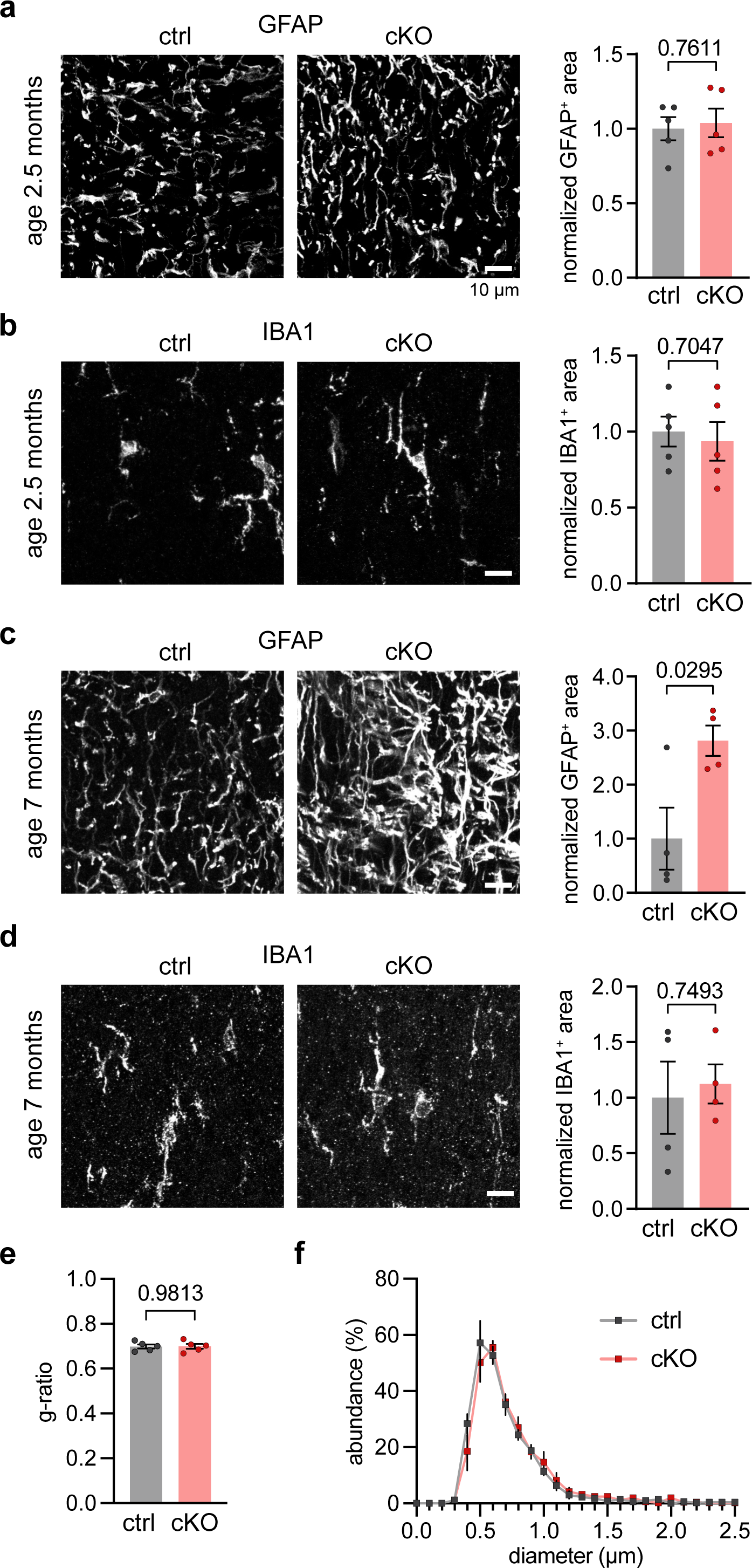
No signs of astrogliosis or microgliosis in young Kir 4.1 cKO mice, but moderate astrogliosis at the age of 7 months. **a** and **b,** Representative confocal images of immunolabeling (**a**) the astrocyte marker GFAP and (**b**) the microglial marker IBA1, from optic nerve sections of 2.5 months old ctrl and cKO mice. At the age of 2.5 months, there were no differences in GFAP labeling (assessed as GFAP-immunopositive area) between the genotypes (n = 4-5 mice, p = 0.7611, unpaired t-test), or in IBA1 immunopositive area (n = 4-5 mice, p = 0.7047, unpaired t-test). **c**, At the age of 7 months, there was a significant increase in GFAP immunopositive area, indicative of astrogliosis, in optic nerves from cKO mice compared to ctrl (n = 4 mice, p = 0.0295, unpaired t-test). **d**, No differences in IBA1 immunopositive area at 7 months of age between the genotypes (n = 4 mice, p = 0.7493, unpaired t-test). **e**, Electron microscopic inspection of optic nerves from 7-8 months old cKO and ctrl mice (see Fig. 3k-m) revealed no difference in myelin sheath thickness determined by g-ratio analysis (n = 5 mice, p = 0.9813, unpaired t-test) and (**f**) normal axon size distribution of myelinated axons (*F_interaction_* (27, 216) = 0.6975, p = 0.8670, two-way ANOVA). Data are represented as means ± SEM.

**Extended Data Figure 6.**
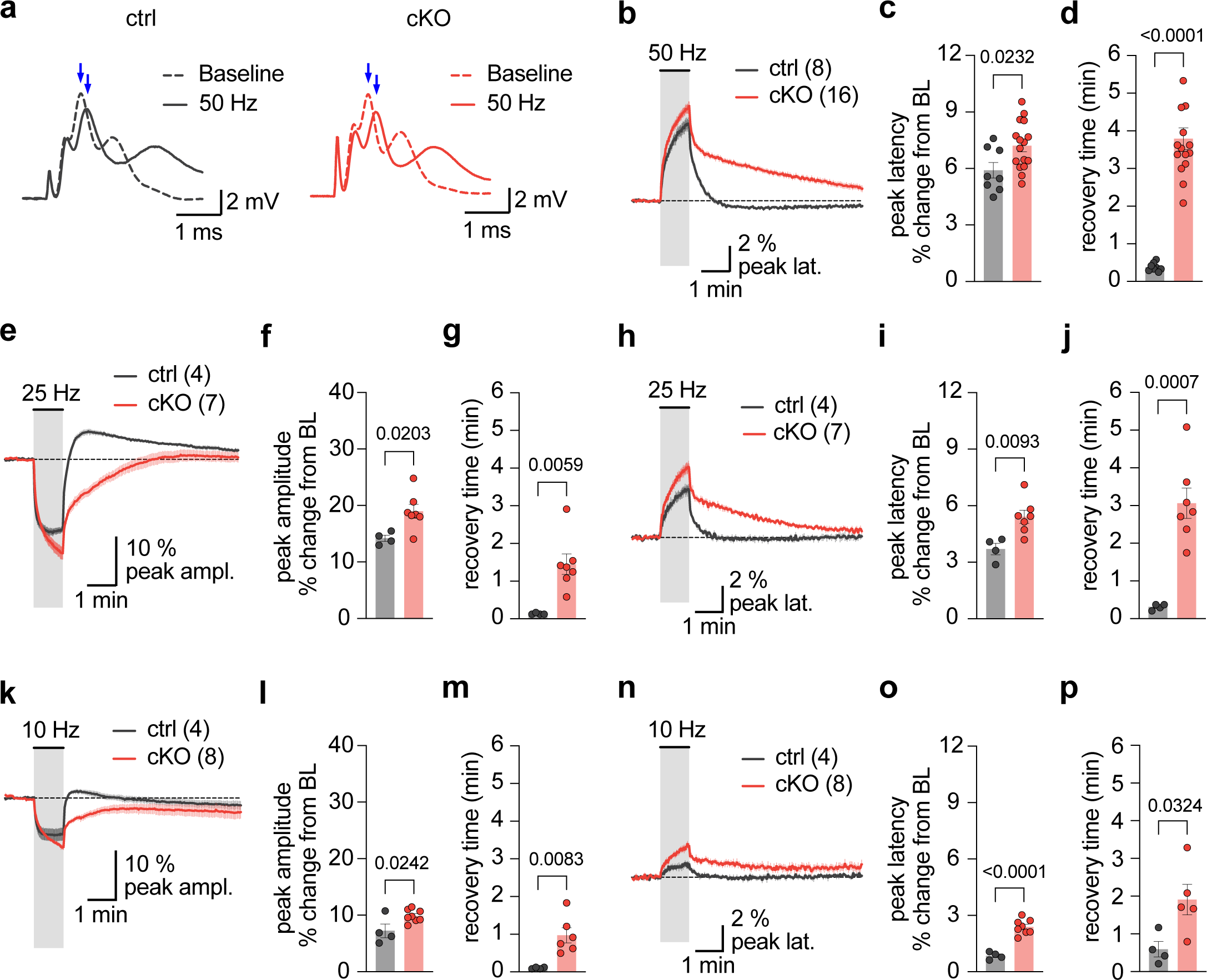
Extended analysis of activity-dependent deficits in CAP peak recovery kinetics in absence of oligodendroglial Kir4.1. **a,** Representative CAP traces of ctrl and cKO at baseline and at the end of the 50 Hz stimulus train. Note the change in CAP peak amplitude and latency (blue arrows). See also Figure 3h-j for CAP peak amplitude analysis. **b - d**, Analysis of CAP peak latency changes in ctrl (n = 8 mice) and cKO (n = 16 mice). **b**, Averaged changes (in % from baseline) of CAP peak latency upon 1 min 50 Hz stimulation. Note that the recovery of the peak latency after stimulation is slower in cKO mice. **c**, Analysis of CAP peak latency change at the end of the 50 Hz stimulation period from recordings in (**b**). The stimulus-evoked increase in latency was larger in cKO compared to ctrol (p = 0.0232, unpaired t-test). **d**, Analysis of the recovery time of CAP peak latency following 50 Hz stimulation. Recovery is significantly slower in cKO compared to ctrl (p < 0.0001, unpaired t-test). **e-j**, Analysis of 25 Hz stimulation-induced changes in (**e-g**) CAP peak amplitude and (**h-j**) CAP peak latency in ctrl (n = 4 mice) and cKO (n = 7 mice). CAP peak amplitude decreases more (**f**) in cKO mice (p = 0.0203, unpaired t-test) and recovers slower (**g**) after stimulation (p = 0.0059). Similarly, CAP peak latency increases more (**i**) in cKO mice (p = 0.0093, unpaired t-test) and recovers slower (**j**) after stimulation (p = 0.0007, unpaired t-test). **k-p**, Analysis of 10 Hz stimulation-induced changes in (**k-m**) CAP peak amplitude and (**n-p**) CAP peak latency in ctrl (n = 4 mice) and cKO (n = 8 mice). CAP peak amplitude decreases more (**i**) in cKO mice (p = 0.0242, unpaired t-test) and recovers slower (**m**) after stimulation (p = 0.0083). Also, CAP peak latency increases more (**o**) in cKO mice (p < 0.0001, unpaired t-test) and recovers slower (**j**) after stimulation (p = 0.0324, unpaired t-test). Data are represented as means ± SEM.

**Extended Data Figure 7.**
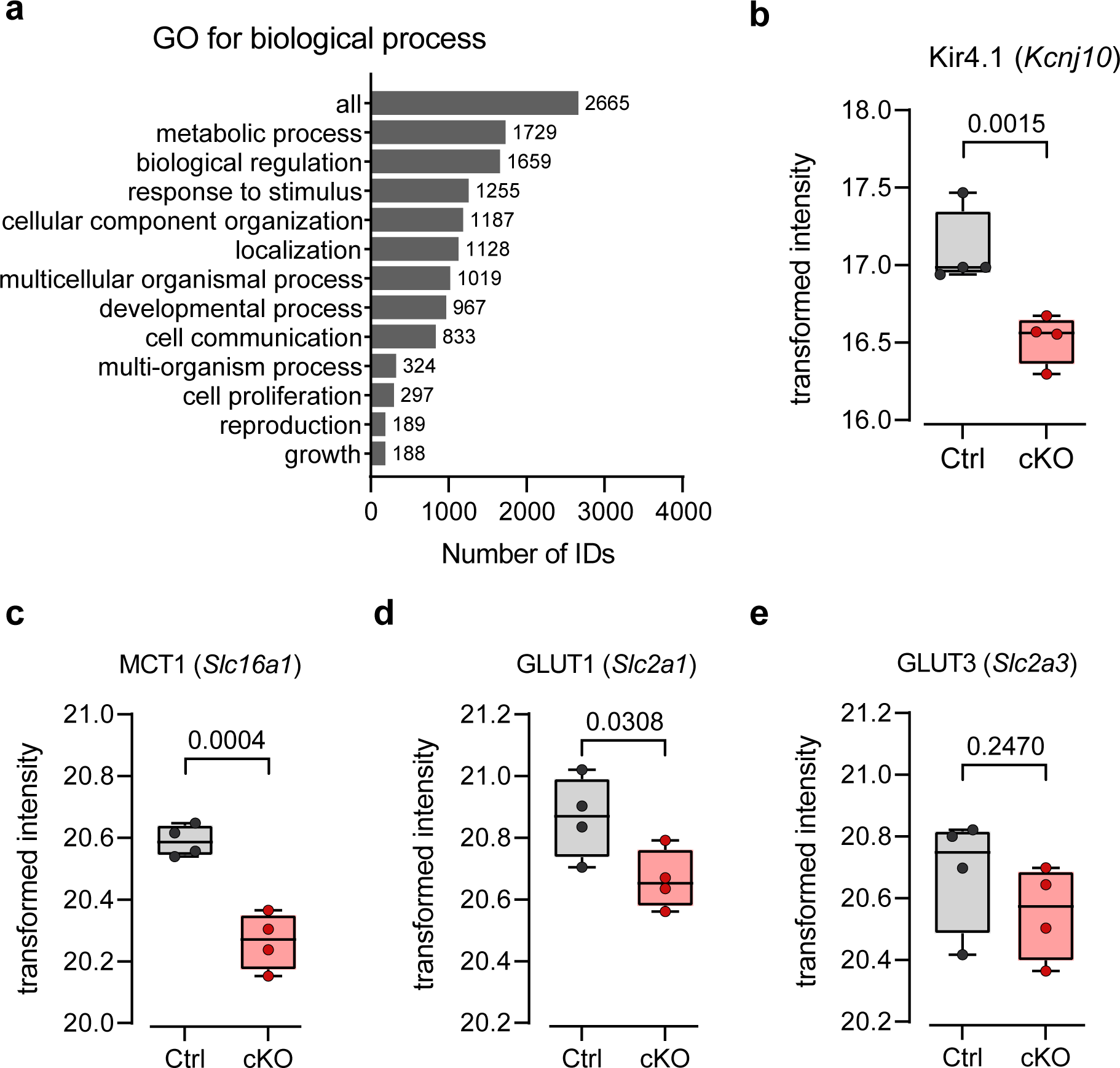
Proteomics analysis revealed reduced GLUT1 and MCT1 abundance in optic nerve lysates from Kir4.1 cKO mice. **a,** Of the 3624 detected protein hits from the TMT-based proteomics analysis, 2665 were unambiguously mapped to unique Entrez gene IDs. Using the GO term biological process, bar charts depict the GO annotation and functional categorization of identified proteins. WebGestalt.org provided the summary. **b**, Protein abundance of Kir4.1 (gene *Kcnj10*) is reduced in samples from cKO (n = 4 mice) compared to ctrl (n = 4, p = 0.0015, moderated t-test). **c**, Abundance of MCT1 (*Slc16a1*) is reduced in cKO compared to ctrl (n = 4, p = 0.0004, moderated t-test). **d**, Abundance of GLUT1 (*Slc2a1*) is reduced in cKO compared to ctrl (n = 4, p = 0.0308, moderated t-test). **e**, Abundance of GLUT3 (*Slc2a3*) is unchanged between genotypes (n = 4, p = 0.2470, moderated t-test). Data are represented as means ± SEM.

**Extended Data Figure 8.**
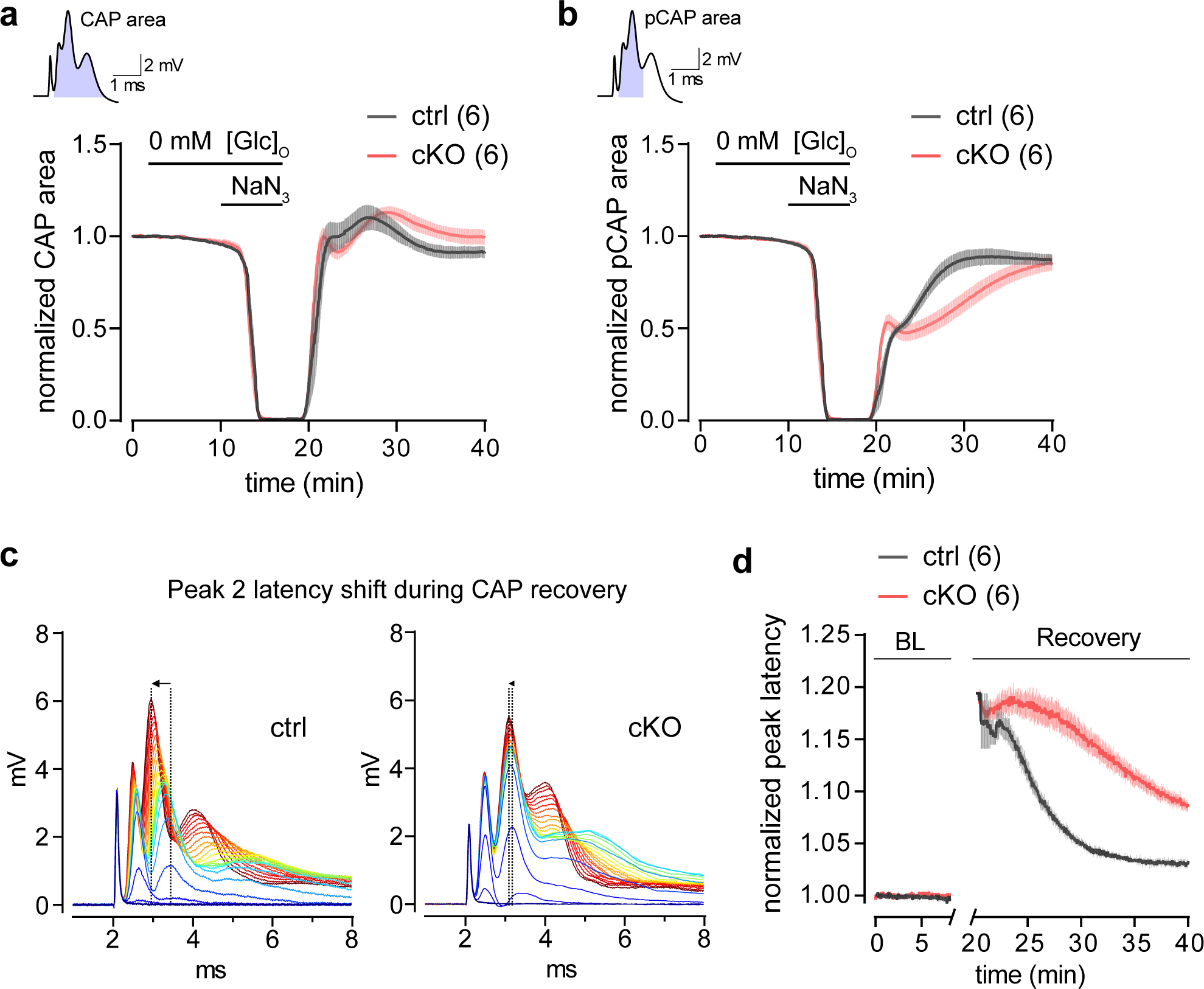
Deficiency in adjusting axonal conduction speed following energy deprivation. **a,** Time course of CAP area (inset, top) changes from optic nerves of ∼3 months old cKO (n = 6) and ctrl mice (n = 6) reversibly challenged with glucose deprivation (GD) and mitochondrial inhibition (MI) with 5 mM NaN_3_ (GD+MI, resembling chemical ischemia). See also Fig. 5b. **b**, Partial CAP (pCAP) area (inset, top) time course analysis during and following GD+MI. Note that the recovery of the pCAP kinetics differs between the genotypes. **d**, Example traces of the gradual recovery of the CAP response (shown in 25 s intervals), shown are the first 10 min of the initial CAP recovery after GD+MI for ctrl (left) and cKO (right). Note that the peak 2 latency shift (arrow at dashed lines) during CAP recovery is larger in ctrl compared to cKO. **e**, Peak 2 latency analysis adjusted to the initial baseline value before GD+MI. Notably, following GD+MI peak 2 latency of the first recovering CAPs is similarly increased in both genotypes, but rapidly returns to normal conduction latency in ctrl, whereas this acceleration of peak latency is much slower in cKO (n = 6, p < 0.0001, two-way ANOVA). Data are represented as means ± SEM.

**Extended Data Figure 9.**
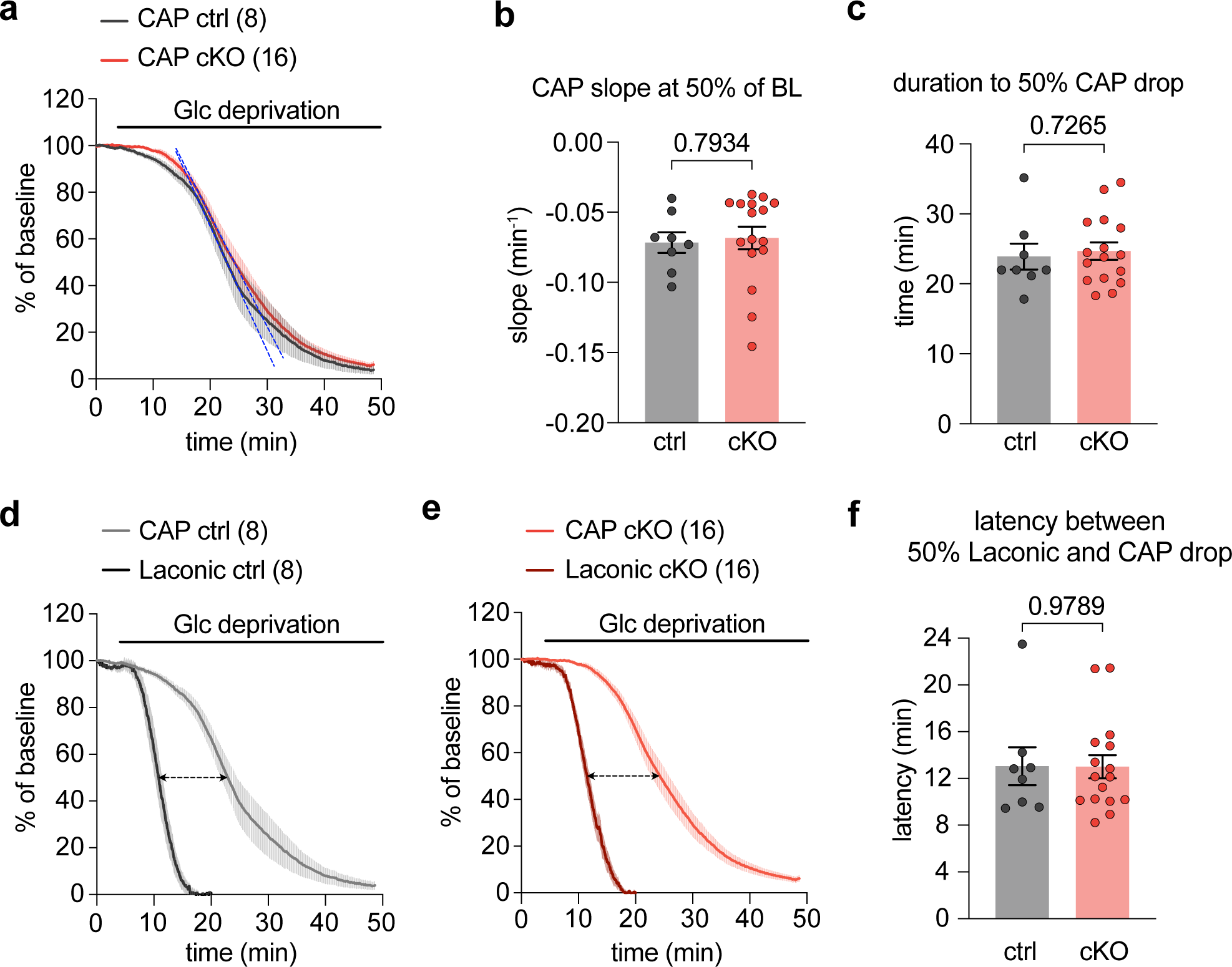
Temporal relationship between axonal lactate and CAP decline during GD. **a-c,** CAP decline kinetics during glucose (Glc) deprivation are comparable between genotypes. **a**, Time course of CAP area decline during glucose deprivation (GD) in ∼3 months old cKO (n = 16) and ctrl mice (n = 8). See also Fig. 5b. **b**, Slope of CAP decline is similar between genotypes (p = 0.7934, unpaired t-test). **c**, Time to 50% CAP drop is comparable between genotypes (p = 0.7265, unpaired t-test). **d** and **e**, CAP and axonal lactate dynamics were simultaneously measured during GD. Depicted are normalized time traces of both recordings for ctrl (**d**) and cKO (**e**). Note that there is a delay between lactate level decline and CAP decline (dashed arrows). Interestingly, the decrease in CAP begins when axonal lactate levels are almost diminished. **f**, The delay between axonal lactate decline and CAP drop is the same between the genotypes (p = 0.9789, unpaired t-test).

**Supplementary Movie 1.** Example time-lapse recording of GCaMP6s-epxressing OLs responding to a 30 s 50 Hz optic nerve stimulation. Images were acquired at 2.96 Hz and movie is played at 160 fps. Stimulation period is indicated by white square (top right corner). Scalebar, 10 µm.

**Supplementary Movie 2.** Example time-lapse recording of GCaMP6s-epxressing optic nerve axons responding to a 30 s 50 Hz optic nerve stimulation. Images were acquired at 2.96 Hz and movie is played at 160 fps. Stimulation period is indicated by white square (top right corner). Scalebar, 10 µm.

## References

1. Salvadores, N., Sanhueza, M., Manque, P. & Court, F. A. Axonal Degeneration during Aging and Its Functional Role in Neurodegenerative Disorders. Front. Neurosci. 11, 451 (2017).

2. Medana, I. M. & Esiri, M. M. Axonal damage: A key predictor of outcome in human CNS diseases. Brain (2003) doi:10.1093/brain/awg061.

3. Tripathi, R. B. et al. Remarkable Stability of Myelinating Oligodendrocytes in Mice. Cell Rep. 21, 316–323 (2017).

4. Saab, A. S., Tzvetanova, I. D. & Nave, K.-A. The role of myelin and oligodendrocytes in axonal energy metabolism. Curr. Opin. Neurobiol. 23, 1065–72 (2013).

5. Nave, K.-A. Myelination and the trophic support of long axons. Nat. Rev. Neurosci. 11, 275–83 (2010).

6. Philips, T. & Rothstein, J. D. Oligodendroglia: metabolic supporters of neurons. J. Clin. Invest. 127, 3271– 3280 (2017).

7. Duncan, G. J., Simkins, T. J. & Emery, B. Neuron-Oligodendrocyte Interactions in the Structure and Integrity of Axons. Front. cell Dev. Biol. 9, 653101 (2021).

8. Xin, W. & Chan, J. R. Myelin plasticity: sculpting circuits in learning and memory. Nat. Rev. Neurosci. 21, 682–694 (2020).

9. Fünfschilling, U. et al. Glycolytic oligodendrocytes maintain myelin and long-term axonal integrity. Nature 485, 517–21 (2012).

10. Lee, Y. et al. Oligodendroglia metabolically support axons and contribute to neurodegeneration. Nature 487, 443–8 (2012).

11. Philips, T. et al. MCT1 Deletion in Oligodendrocyte Lineage Cells Causes Late-Onset Hypomyelination and Axonal Degeneration. Cell Rep. 34, 108610 (2021).

12. Saab, A. S. et al. Oligodendroglial NMDA Receptors Regulate Glucose Import and Axonal Energy Metabolism. Neuron 91, 119–32 (2016).

13. Edgar, J. M. et al. Río-Hortega’s drawings revisited with fluorescent protein defines a cytoplasm-filled channel system of CNS myelin. J. Anat. 239, 1241–1255 (2021).

14. Saab, A. S. & Nave, K.-A. Myelin dynamics: protecting and shaping neuronal functions. Curr. Opin. Neurobiol. 47, 104–112 (2017).

15. Snaidero, N. et al. Antagonistic Functions of MBP and CNP Establish Cytosolic Channels in CNS Myelin. Cell Rep. 18, 314–323 (2017).

16. Trevisiol, A. et al. Structural myelin defects are associated with low axonal ATP levels but rapid recovery from energy deprivation in a mouse model of spastic paraplegia. PLoS Biol. 18, e3000943 (2020).

17. Griffiths, I. et al. Axonal swellings and degeneration in mice lacking the major proteolipid of myelin. Science 280, 1610–3 (1998).

18. Lüders, K. A. et al. Maintenance of high proteolipid protein level in adult central nervous system myelin is required to preserve the integrity of myelin and axons. Glia 67, 634–649 (2019).

19. Edgar, J. M. et al. Oligodendroglial modulation of fast axonal transport in a mouse model of hereditary spastic paraplegia. J. Cell Biol. (2004) doi:10.1083/jcb.200312012.

20. Steyer, A. M. et al. Pathology of myelinated axons in the PLP-deficient mouse model of spastic paraplegia type 2 revealed by volume imaging using focused ion beam-scanning electron microscopy. J. Struct. Biol. 210, 107492 (2020).

21. Mukherjee, C. et al. Oligodendrocytes Provide Antioxidant Defense Function for Neurons by Secreting Ferritin Heavy Chain. Cell Metab. 32, 259–272.e10 (2020).

22. Larson, V. A. et al. Oligodendrocytes control potassium accumulation in white matter and seizure susceptibility. Elife 7, (2018).

23. Schirmer, L. et al. Oligodendrocyte-encoded Kir4.1 function is required for axonal integrity. Elife 7, (2018).

24. Looser, Z. J., Barrett, M. J. P., Hirrlinger, J., Weber, B. & Saab, A. S. Intravitreal AAV-Delivery of Genetically Encoded Sensors Enabling Simultaneous Two-Photon Imaging and Electrophysiology of Optic Nerve Axons. Front. Cell. Neurosci. 12, 377 (2018).

25. Trevisiol, A. et al. Monitoring ATP dynamics in electrically active white matter tracts. Elife 6, (2017).

26. Doerflinger, N. H., Macklin, W. B. & Popko, B. Inducible site-specific recombination in myelinating cells. Genesis 35, 63–72 (2003).

27. Chen, T.-W. et al. Ultrasensitive fluorescent proteins for imaging neuronal activity. Nature 499, 295–300 (2013).

28. Madisen, L. et al. Transgenic mice for intersectional targeting of neural sensors and effectors with high specificity and performance. Neuron 85, 942–58 (2015).

29. Micu, I. et al. The molecular physiology of the axo-myelinic synapse. Exp. Neurol. 276, 41–50 (2016).

30. Micu, I. et al. NMDA receptors mediate calcium accumulation in myelin during chemical ischaemia. Nature 439, 988–92 (2006).

31. James, G. & Butt, A. M. P2X and P2Y purinoreceptors mediate ATP-evoked calcium signalling in optic nerve glia in situ. Cell Calcium 30, 251–9 (2001).

32. Kirischuk, S., Scherer, J., Kettenmann, H. & Verkhratsky, A. Activation of P2-purinoreceptors triggered Ca2+ release from InsP3-sensitive internal stores in mammalian oligodendrocytes. J. Physiol. 483 (**Pt 1**), 41–57 (1995).

33. Matute, C. et al. P2X(7) receptor blockade prevents ATP excitotoxicity in oligodendrocytes and ameliorates experimental autoimmune encephalomyelitis. J.Neurosci. 27, 9525–9533 (2007).

34. Stevens, B., Porta, S., Haak, L. L., Gallo, V. & Fields, R. D. Adenosine: a neuron-glial transmitter promoting myelination in the CNS in response to action potentials. Neuron 36, 855–68 (2002).

35. Ransom, C. B., Ransom, B. R. & Sontheimer, H. Activity-dependent extracellular K+ accumulation in rat optic nerve: the role of glial and axonal Na+ pumps. J. Physiol. 522 **Pt** **3**, 427–42 (2000).

36. Bay, V. & Butt, A. M. Relationship between glial potassium regulation and axon excitability: a role for glial Kir4.1 channels. Glia 60, 651–60 (2012).

37. Olsen, M. L. & Sontheimer, H. Functional implications for Kir4.1 channels in glial biology: from K+ buffering to cell differentiation. J. Neurochem. 107, 589–601 (2008).

38. Yamazaki, Y. et al. Modulatory effects of oligodendrocytes on the conduction velocity of action potentials along axons in the alveus of the rat hippocampal CA1 region. Neuron Glia Biol. (2007) doi:10.1017/S1740925X08000070.

39. Battefeld, A., Klooster, J. & Kole, M. H. P. Myelinating satellite oligodendrocytes are integrated in a glial syncytium constraining neuronal high-frequency activity. Nat. Commun. 7, 11298 (2016).

40. Boscia, F. et al. Silencing or knocking out the Na(+)/Ca(2+) exchanger-3 (NCX3) impairs oligodendrocyte differentiation. Cell Death Differ. 19, 562–72 (2012).

41. Casamassa, A. et al. Ncx3 gene ablation impairs oligodendrocyte precursor response and increases susceptibility to experimental autoimmune encephalomyelitis. Glia 64, 1124–37 (2016).

42. Spencer, S. A., Suárez-Pozos, E., Escalante, M., Myo, Y. P. & Fuss, B. Sodium-Calcium Exchangers of the SLC8 Family in Oligodendrocytes: Functional Properties in Health and Disease. Neurochem. Res. 45, 1287– 1297 (2020).

43. Friess, M. et al. Intracellular ion signaling influences myelin basic protein synthesis in oligodendrocyte precursor cells. Cell Calcium 60, 322–330 (2016).

44. Moyon, S. et al. TET1-mediated DNA hydroxymethylation regulates adult remyelination in mice. Nat. Commun. 12, 3359 (2021).

45. Yamazaki, Y., Abe, Y., Fujii, S. & Tanaka, K. F. Oligodendrocytic Na+-K+-Cl-co-transporter 1 activity facilitates axonal conduction and restores plasticity in the adult mouse brain. Nat. Commun. 12, 5146 (2021).

46. Djukic, B., Casper, K. B., Philpot, B. D., Chin, L.-S. & McCarthy, K. D. Conditional knock-out of Kir4.1 leads to glial membrane depolarization, inhibition of potassium and glutamate uptake, and enhanced short-term synaptic potentiation. J. Neurosci. 27, 11354–65 (2007).

47. Imamura, H. et al. Visualization of ATP levels inside single living cells with fluorescence resonance energy transfer-based genetically encoded indicators. Proc. Natl. Acad. Sci. U. S. A. 106, 15651–6 (2009).

48. Hamilton, N. B., Kolodziejczyk, K., Kougioumtzidou, E. & Attwell, D. Proton-gated Ca(2+)-permeable TRP channels damage myelin in conditions mimicking ischaemia. Nature 529, 523–7 (2016).

49. San Martín, A., et al. A genetically encoded FRET lactate sensor and its use to detect the Warburg effect in single cancer cells. PLoS One 8, e57712 (2013).

50. Meyer, N. et al. Oligodendrocytes in the Mouse Corpus Callosum Maintain Axonal Function by Delivery of Glucose. Cell Rep 22, 2383–2394 (2018).

51. Takanaga, H., Chaudhuri, B. & Frommer, W. B. GLUT1 and GLUT9 as major contributors to glucose influx in HepG2 cells identified by a high sensitivity intramolecular FRET glucose sensor. Biochim. Biophys. Acta 1778, 1091–9 (2008).

52. Bittner, C. X. et al. High resolution measurement of the glycolytic rate. Front. Neuroenergetics 2, (2010).

53. Bittner, C. X. et al. Fast and reversible stimulation of astrocytic glycolysis by K+ and a delayed and persistent effect of glutamate. J. Neurosci. 31, 4709–13 (2011).

54. Zhang, X. et al. Oligodendroglial glycolytic stress triggers inflammasome activation and neuropathology in Alzheimer’s disease. Sci. Adv. 6, (2020).

55. Mot, A. I., Depp, C. & Nave, K.-A. An emerging role of dysfunctional axon-oligodendrocyte coupling in neurodegenerative diseases. Dialogues Clin. Neurosci. 20, 283–292 (2018).

56. Kenigsbuch, M. et al. A shared disease-associated oligodendrocyte signature among multiple CNS pathologies. Nat. Neurosci. 25, 876–886 (2022).

57. Kaya, T. et al. CD8+ T cells induce interferon-responsive oligodendrocytes and microglia in white matter aging. Nat. Neurosci. (2022) doi:10.1038/s41593-022-01183-6.

58. Kettenmann, H., Sonnhof, U. & Schachner, M. Exclusive potassium dependence of the membrane potential in cultured mouse oligodendrocytes. J. Neurosci. 3, 500–5 (1983).

59. Brasko, C., Hawkins, V., De La Rocha, I. C. & Butt, A. M. Expression of Kir4.1 and Kir5.1 inwardly rectifying potassium channels in oligodendrocytes, the myelinating cells of the CNS. Brain Struct. Funct. 222, 41–59 (2017).

60. Papanikolaou, M., Butt, A. M. & Lewis, A. A critical role for the inward rectifying potassium channel Kir7.1 in oligodendrocytes of the mouse optic nerve. Brain Struct. Funct. (2020) doi:10.1007/s00429-020-02043-4.

61. Micu, I., Plemel, J. R., Caprariello, A. V, Nave, K. A. & Stys, P. K. Axo-myelinic neurotransmission: a novel mode of cell signalling in the central nervous system. Nat Rev Neurosci 19, 49–58 (2018).

62. Almeida, R. G. et al. Myelination induces axonal hotspots of synaptic vesicle fusion that promote sheath growth. Curr. Biol. 31, 3743–3754.e5 (2021).

63. Hines, J. H., Ravanelli, A. M., Schwindt, R., Scott, E. K. & Appel, B. Neuronal activity biases axon selection for myelination in vivo. Nat. Neurosci. 18, 683–9 (2015).

64. Wake, H., Lee, P. R. & Fields, R. D. Control of local protein synthesis and initial events in myelination by action potentials. Science 333, 1647–51 (2011).

65. Mensch, S. et al. Synaptic vesicle release regulates myelin sheath number of individual oligodendrocytes in vivo. Nat. Neurosci. 18, 628–30 (2015).

66. Kukley, M., Capetillo-Zarate, E. & Dietrich, D. Vesicular glutamate release from axons in white matter. Nat. Neurosci. 10, 311–20 (2007).

67. Ziskin, J. L., Nishiyama, A., Rubio, M., Fukaya, M. & Bergles, D. E. Vesicular release of glutamate from unmyelinated axons in white matter. Nat. Neurosci. 10, 321–30 (2007).

68. Krasnow, A. M., Ford, M. C., Valdivia, L. E., Wilson, S. W. & Attwell, D. Regulation of developing myelin sheath elongation by oligodendrocyte calcium transients in vivo. Nat. Neurosci. 21, 24–28 (2018).

69. Baraban, M., Koudelka, S. & Lyons, D. A. Ca 2+ activity signatures of myelin sheath formation and growth in vivo. Nat. Neurosci. 21, 19–23 (2018).

70. Battefeld, A., Popovic, M. A., de Vries, S. I. & Kole, M. H. P. High-Frequency Microdomain Ca2+ Transients and Waves during Early Myelin Internode Remodeling. Cell Rep. 26, 182–191.e5 (2019).

71. Kanda, H. et al. TREK-1 and TRAAK Are Principal K+ Channels at the Nodes of Ranvier for Rapid Action Potential Conduction on Mammalian Myelinated Afferent Nerves. Neuron 104, 960–971.e7 (2019).

72. Brohawn, S. G. et al. The mechanosensitive ion channel TRAAK is localized to the mammalian node of Ranvier. Elife 8, (2019).

73. Rash, J. E. Molecular disruptions of the panglial syncytium block potassium siphoning and axonal saltatory conduction: pertinence to neuromyelitis optica and other demyelinating diseases of the central nervous system. Neuroscience 168, 982–1008 (2010).

74. Cohen, C. C. H. et al. Saltatory Conduction along Myelinated Axons Involves a Periaxonal Nanocircuit. Cell 180, 311–322.e15 (2020).

75. Menichella, D. M. et al. Genetic and physiological evidence that oligodendrocyte gap junctions contribute to spatial buffering of potassium released during neuronal activity. J. Neurosci. 26, 10984–91 (2006).

76. Neusch, C., Rozengurt, N., Jacobs, R. E., Lester, H. A. & Kofuji, P. Kir4.1 potassium channel subunit is crucial for oligodendrocyte development and in vivo myelination. J. Neurosci. 21, 5429–38 (2001).

77. Armbruster, M. et al. Neuronal activity drives pathway-specific depolarization of peripheral astrocyte processes. Nat. Neurosci. 25, 607–616 (2022).

78. Orthmann-Murphy, J. L., Abrams, C. K. & Scherer, S. S. Gap junctions couple astrocytes and oligodendrocytes. J. Mol. Neurosci. 35, 101–16 (2008).

79. Hösli, L. et al. Decoupling astrocytes in adult mice impairs synaptic plasticity and spatial learning. Cell Rep. 38, 110484 (2022).

80. Fernández-Moncada, I. et al. Bidirectional astrocytic GLUT1 activation by elevated extracellular K. Glia 69, 1012–1021 (2021).

81. Ruminot, I., Schmälzle, J., Leyton, B., Barros, L. F. & Deitmer, J. W. Tight coupling of astrocyte energy metabolism to synaptic activity revealed by genetically encoded FRET nanosensors in hippocampal tissue. J. Cereb. Blood Flow Metab. 39, 513–523 (2019).

82. Sotelo-Hitschfeld, T. et al. Channel-mediated lactate release by K^+^-stimulated astrocytes. J. Neurosci. 35, 4168–78 (2015).

83. Zuend, M. et al. Arousal-induced cortical activity triggers lactate release from astrocytes. Nat. Metab. (2020) doi:10.1038/s42255-020-0170-4.

84. Jahn, O. et al. The CNS Myelin Proteome: Deep Profile and Persistence After Post-mortem Delay. Front. Cell. Neurosci. 14, 239 (2020).

85. Zhang, Y. et al. An RNA-sequencing transcriptome and splicing database of glia, neurons, and vascular cells of the cerebral cortex. J. Neurosci. 34, 11929–47 (2014).

86. Gargareta, V.-I. et al. Conservation and divergence of myelin proteome and oligodendrocyte transcriptome profiles between humans and mice. Elife 11, (2022).

87. Leto, D. & Saltiel, A. R. Regulation of glucose transport by insulin: traffic control of GLUT4. Nat. Rev. Mol. Cell Biol. 13, 383–96 (2012).

88. Martin, L. B., Shewan, A., Millar, C. A., Gould, G. W. & James, D. E. Vesicle-associated membrane protein 2 plays a specific role in the insulin-dependent trafficking of the facilitative glucose transporter GLUT4 in 3T3-L1 adipocytes. J. Biol. Chem. 273, 1444–52 (1998).

89. Lodhi, I. J. et al. Gapex-5, a Rab31 guanine nucleotide exchange factor that regulates Glut4 trafficking in adipocytes. Cell Metab. 5, 59–72 (2007).

90. Lam, M. et al. CNS myelination requires VAMP2/3-mediated membrane expansion in oligodendrocytes. Nat. Commun. 13, 5583 (2022).

91. Feldmann, A. et al. Transport of the major myelin proteolipid protein is directed by VAMP3 and VAMP7. J. Neurosci. 31, 5659–72 (2011).

92. White, R. & Krämer-Albers, E.-M. Axon-glia interaction and membrane traffic in myelin formation. Front. Cell. Neurosci. 7, (2014).

93. Dienel, G. A. Brain Glucose Metabolism: Integration of Energetics with Function. Physiol. Rev. 99, 949– 1045 (2019).

94. Barros, L. F. et al. Fluid Brain Glycolysis: Limits, Speed, Location, Moonlighting, and the Fates of Glycogen and Lactate. Neurochem. Res. 45, 1328–1334 (2020).

95. Herrero-Mendez, A. et al. The bioenergetic and antioxidant status of neurons is controlled by continuous degradation of a key glycolytic enzyme by APC/C-Cdh1. Nat. Cell Biol. 11, 747–52 (2009).

96. Zala, D. et al. Vesicular glycolysis provides on-board energy for fast axonal transport. Cell 152, 479–91 (2013).

97. Frühbeis, C. et al. Oligodendrocytes support axonal transport and maintenance via exosome secretion. PLoS Biol. 18, e3000621 (2020).

98. Chamberlain, K. A. et al. Oligodendrocytes enhance axonal energy metabolism by deacetylation of mitochondrial proteins through transcellular delivery of SIRT2. Neuron 109, 3456–3472.e8 (2021).

99. Frühbeis, C. et al. Neurotransmitter-triggered transfer of exosomes mediates oligodendrocyte-neuron communication. PLoS Biol. 11, e1001604 (2013).

100. Baeza-Lehnert, F. et al. Non-Canonical Control of Neuronal Energy Status by the Na(+) Pump. Cell Metab (2018) doi:10.1016/j.cmet.2018.11.005.

101. Hövelmeyer, N. et al. Apoptosis of oligodendrocytes via Fas and TNF-R1 is a key event in the induction of experimental autoimmune encephalomyelitis. J. Immunol. 175, 5875–84 (2005).

102. Fleischmann, T., Jirkof, P., Henke, J., Arras, M. & Cesarovic, N. Injection anaesthesia with fentanyl-midazolam-medetomidine in adult female mice: importance of antagonization and perioperative care. Lab. Anim. 50, 264–74 (2016).

103. Paterna, J.-C., Feldon, J. & Büeler, H. Transduction profiles of recombinant adeno-associated virus vectors derived from serotypes 2 and 5 in the nigrostriatal system of rats. J. Virol. 78, 6808–17 (2004).

104. Mayrhofer, J. M. et al. Design and performance of an ultra-flexible two-photon microscope for in vivo research. Biomed. Opt. Express 6, 4228–37 (2015).

105. Pologruto, T. A., Sabatini, B. L. & Svoboda, K. ScanImage: flexible software for operating laser scanning microscopes. Biomed Eng Online 2, 13 (2003).

106. Stys, P. K., Ransom, B. R. & Waxman, S. G. Compound action potential of nerve recorded by suction electrode: a theoretical and experimental analysis. Brain Res. 546, 18–32 (1991).

107. Barrett, M. J. P., Ferrari, K. D., Stobart, J. L., Holub, M. & Weber, B. CHIPS: an Extensible Toolbox for Cellular and Hemodynamic Two-Photon Image Analysis. Neuroinformatics 16, 145–147 (2018).

108. Glück, C. et al. Distinct signatures of calcium activity in brain mural cells. Elife 10, (2021).

109. Möbius, W. et al. Electron microscopy of the mouse central nervous system. Methods Cell Biol. 96, 475– 512 (2010).

110. Türker, C. et al. B-Fabric. in Proceedings of the 13th International Conference on Extending Database Technology - EDBT ’10 717 (ACM Press, 2010). doi:10.1145/1739041.1739135.

111. Wolski, W., Panse, C., Grossmann, J., D’Errico, M. & Nanni, P. prolfqua - R-package for Proteomics Label Free Quantification using Linear Models Implementation & Methods Benchmarking References. in *poster*, F1000Research 2021 (2021). doi:https://doi.org/10.7490/f1000research.1118455.1.

112. Ritchie, M. E. et al. limma powers differential expression analyses for RNA-sequencing and microarray studies. Nucleic Acids Res. 43, e47 (2015).

113. Liao, Y., Wang, J., Jaehnig, E. J., Shi, Z. & Zhang, B. WebGestalt 2019: gene set analysis toolkit with revamped UIs and APIs. Nucleic Acids Res. 47, W199–W205 (2019).

114. Erwig, M. S. et al. Myelin: Methods for Purification and Proteome Analysis. Methods Mol. Biol. 1936, 37– 63 (2019).

115. Stumpf, S. K. et al. Ketogenic diet ameliorates axonal defects and promotes myelination in Pelizaeus-Merzbacher disease. Acta Neuropathol. 138, 147–161 (2019).

116. Berghoff, S. A. et al. Blood-brain barrier hyperpermeability precedes demyelination in the cuprizone model. Acta Neuropathol. Commun. 5, 94 (2017).

117. Jung, M., Sommer, I., Schachner, M. & Nave, K. A. Monoclonal antibody O10 defines a conformationally sensitive cell-surface epitope of proteolipid protein (PLP): evidence that PLP misfolding underlies dysmyelination in mutant mice. J. Neurosci. 16, 7920–9 (1996).

